# A metabolic coincidence mechanism controls winter photoperiodism in plants

**DOI:** 10.1101/2020.11.13.381426

**Authors:** Wei Liu, Ann Feke, Chun Chung Leung, Daniel A. Tarté, Wenxin Yuan, Morgan Vanderwall, Garrett Sager, Xing Wu, Ariela Schear, Damon A. Clark, Bryan C. Thines, Joshua M. Gendron

## Abstract

Plants have served as a preeminent study system for photoperiodism because of their propensity to flower in concordance with the seasons. A nearly singular focus on understanding seasonal flowering has been to the detriment of discovering other photoperiod measuring mechanisms that may be necessary for vegetative health. Here we use bioinformatics to identify a group of winter photoperiod-induced genes in Arabidopsis and show that one, *PP2-A13*, is critical for fitness and survival, exclusively in winter-like photoperiods. We create a real-time photoperiod reporter, using the *PP2-A13* promoter driving luciferase, and show that winter photoperiod genes are regulated independent of the canonical CO/FT mechanism for photoperiodic flowering. The reporter then allows us to identify the first genetic and cellular drivers of winter photoperiodism and reveal a mechanism that relies on coincidence between light capture through photosynthesis and rhythmic metabolism. This work demonstrates that plants have distinct photoperiod measuring mechanisms that enact critical biological and developmental processes in different seasons.

## Introduction

The obliquity of Earth results in day and night durations (photoperiods) that change throughout the year in most places on earth. Photoperiod is a highly predictable environmental signal that can help organisms anticipate impending seasonal changes (Nelson, et al., 2010). Photoperiod measuring mechanisms are found in fungi (Tan, et al., 2004; Roenneberg and Merrow, 2001), plants (Shim and Imaizumi, 2015; Song, et al., 2015), and animals (Saunders, 2020; Nakane and Yoshimura, 2019) and coordinate seasonal developmental programs to mitigate damage from less predictable abiotic and biotic stresses (Walker, et al., 2019). They also act to align growth and reproduction with seasons that are optimal for organismal fitness. Furthermore, human syndromes, such as seasonal affective disorder and its comorbidities, are under the control of photoperiod (Garbazza and Benedetti, 2018).

Plants have long been a preeminent study system for understanding photoperiod measurement mechanisms because flowering time is easily observable and is often regulated by photoperiod. In the early part of the 20^th^ century, Erwin Bünning used flowering time studies to postulate a two state model for photoperiod measuring systems (Saunders, 2005; Bunning, 1969). In the first part of the 24-hour day, organisms are in a photophilic (light-loving) state and then later in the day they switch to a skotophilic (dark-loving) state. Bünning postulated that a circadian clock controls the phasing of the photophilic and skotophilic states during the 24-hour day. This underlying two-state mechanism allows the organism to enact different developmental programs depending on whether dusk coincides with the photophilic or skotophilic state. For instance, winter dusk occurs in the photophilic state, or early day state, and the organism has one developmental outcome (i.e. vegetative growth in a “long day” flowering plant). Conversely, summer dusk occurs in the skotophilic state, or late day state, and a different outcome occurs (i.e. flowering in a “long day” flowering plant). These criteria allow for a so-called “true photoperiod measuring mechanism” that counts the number of hours of light or dark each day, irrespective of light intensity.

With seasonal flowering, Bünning’s century-old theory held true. Photoperiodic time measurement in flowering relies on circadian clock-controlled transcription of a gene called *CONSTANS* (*CO*) (Putterill, et al., 1995). In Arabidopsis, *CO* mRNA expression is phased to the latter (skotophilic) portion of the 24-hour day, thus low and high *CO* mRNA levels define the photophilic and skotophilic states, respectively (Yanovsky and Kay, 2002). Photoperiodic time measurement then occurs through light-mediated stabilization of CO protein when day length is extended into the skotophilic phase, the time when *CO* mRNA levels are high (Jang, et al., 2008). CO protein subsequently activates transcription of *FLOWERING LOCUS T* (*FT*) that encodes the tissue-mobile florigen (An, et al., 2004; Valverde, et al., 2004; Kardailsky, et al., 1999).

While studies in Arabidopsis have generated immense knowledge of the molecular determinants for photoperiod-controlled flowering, far less is known about other photoperiod-controlled processes in plants. This is especially true for the physiological and cellular processes that are induced in winter-like photoperiods to maintain cellular health. Along with lower average temperatures and changes in water availability, winter poses a unique challenge for plants due to the lower average amount of light that can be used for energy production (Vitasse, et al., 2014; Oquist and Huner, 2003). Despite the potential danger, winter is also necessary for survival in many plants and provides them with a yearly “memory” to distinguish between identical photoperiods throughout the year (Bouche, et al., 2017; Henderson, et al., 2003). Currently, perennial trees have served as models for winter photoperiod-induced dormancy and growth cessation, and recent technological advances have allowed researchers to predict that a variation of the CO/FT module used for flowering is likely playing a role in repression of winter photoperiod transcripts in long summer-like days (Cubas, 2020; Azeez and Sane, 2015; Bohlenius, et al., 2006). However, the gene regulatory networks that control induction of winter photoperiod transcripts have not been studied in detail, and it has been postulated that winter photoperiod induced biological processes could simply be activated by the absence of summer repressive mechanisms. Alternatively, it is possible that there is a wholly separate winter photoperiod transcript induction mechanism. It is likely that we have yet to make this distinction due to a lack of tools to study winter photoperiod processes and sparse knowledge of the genes and cellular processes that are induced in plants in winter photoperiods.

To address this gap, we analyzed genome-wide expression data using daily expression integral calculations to identify transcripts whose expression are induced in winter-like photoperiods in Arabidopsis. Strikingly, we found one prevailing dark biphasic expression pattern associated with transcripts that are induced by winter photoperiods. We characterized the function of one winter photoperiod-induced gene, *PHLOEM PROTEIN2-A13 (PP2-A13)*, showing that it is necessary for cellular health and reproduction in winter-like photoperiods and controls glycoprotein abundance and functions in parallel to autophagy in plants. We created a *PP2-A13*_*promoter*_::*luciferase* transgenic plant, that acts as a real-time photoperiod reporter, and define the properties of the winter transcript induction system demonstrating that it is independent of the CO/FT photoperiod measuring system. We then show that the system relies on light sensing by photosynthesis and that darkness is interpreted by a mechanism that is controlled by rhythmic metabolism. Together, these results show that a metabolic coincidence mechanism drives winter photoperiod transcript induction and define a new photoperiod measuring system that is critical for cellular and physiological health in plants growing in winter photoperiods.

## Results

### Calculating relative daily expression integrals to identify photoperiod-induced transcripts and biological processes

The well-studied photoperiod-induced flowering time gene, *FT*, has a daily expression rhythm in Arabidopsis with high amplitude in 16 hours light:8 hours dark (16L:8D) growth conditions, and low or no amplitude in 8 hours light:16 hours dark (8L:16D) (Yanovsky and Kay, 2002; Suarez-Lopez, et al., 2001). We surmised that other photoperiod-induced transcripts may also be identified through a photoperiod-specific daily rhythm. We estimated daily expression induction by calculating a relative daily expression integral (rDEI = sum of 24 hours expression in condition one / sum of 24 hours expression in condition two) (Figure 1A). To find transcripts that were induced in long summer-like days and short winter-like days we calculated a rDEI using gene expression data from plants grown in 8 hours light:16 hours dark (8L:16D) or 16 hours light:8 hours dark (16L:8D) day growth conditions (rDEI_8L:16D/16L:8D_) (Figure 1A and Table S1) (Michael, et al., 2008; Mockler, et al., 2007). According to our calculations, 359 transcripts are induced greater than two-fold in plants grown in an 8L:16D photoperiod, and 194 transcripts are induced greater than two-fold in plants grown in a 16L:8D photoperiod. Clustering analyses revealed 4 co-expression clusters in the 8L:16D-induced transcripts and 4 clusters in the 16L:8D-induced transcripts (Figure 1B, S1 and Table S2, S3). Approximately 88% of the transcripts with rDEI_8L:16D/16L:8D_ > 2.0 are phased to the dark part of the photoperiod, suggesting that nighttime expression is important for an 8L:16D-induced gene expression signature (316/359; 8L:16D clusters A_W_-C_W_; Fig.1B). Conversely, 73% of the 16L:8D-induced transcripts were phased to the light part of the photoperiod (141/194; 16L:8D Clusters A_S_ and B_S_; Figure S1).

**Figure 1.**
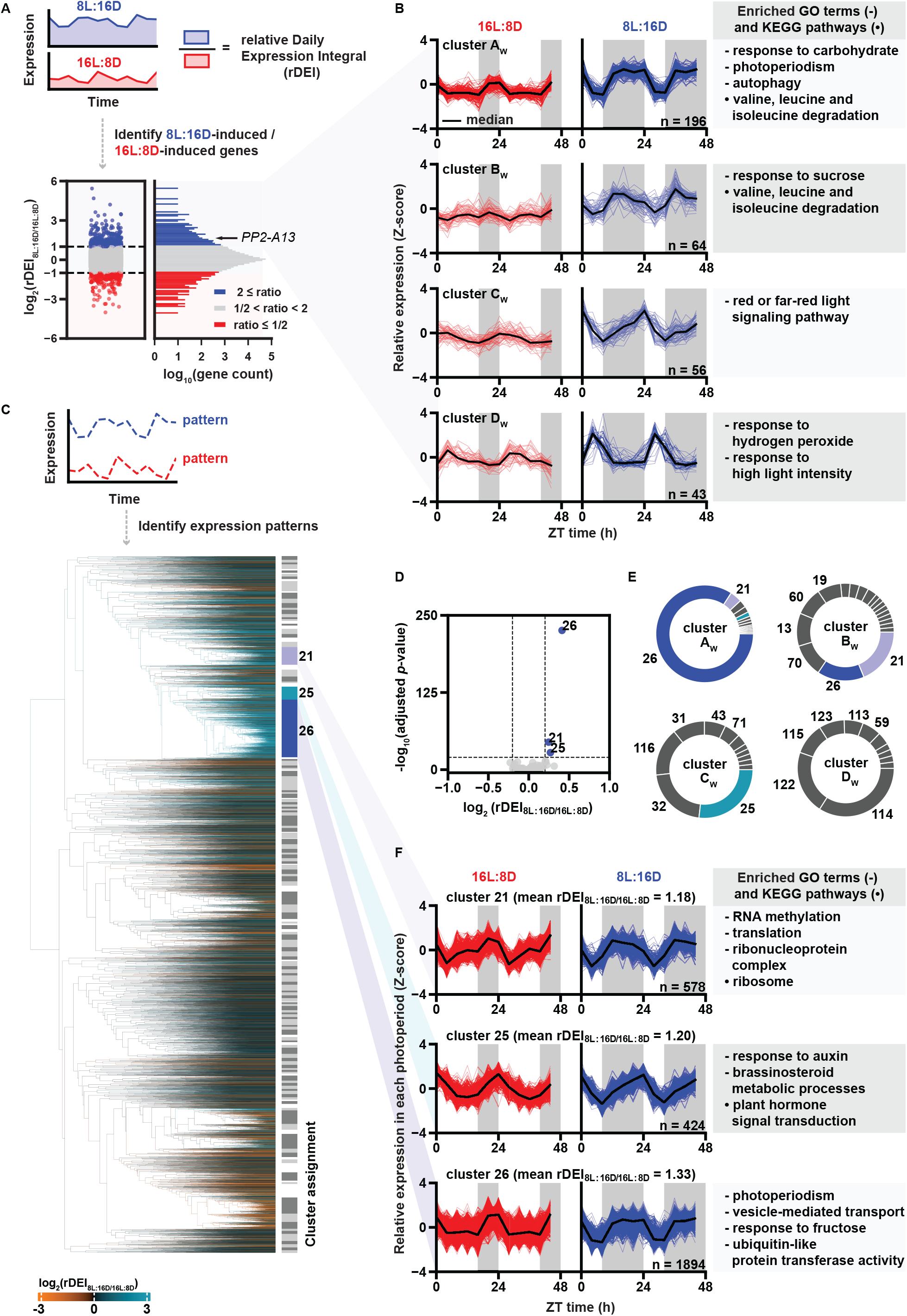
Induced gene expression in an 8L:16D photoperiod is correlated to rhythmic expression patterns with nighttime phasing. (A) Identification of photoperiod-induced transcripts from the DIURNAL Affymetrix ATH1 microarray dataset using relative daily expression integral (rDEI; ratio of sum of expression between two time courses). Distribution of transcript rDEI_8L:16D/16L:8D_ are presented in the histogram (n = 22810). Blue: 8L:16D-induced transcripts with rDEI_8L:16D/16L:8D_ > 2.0; red: 16L:8D-induced transcripts with rDEI_8L:16D/16L:8D_ < 0.5; grey, all other transcripts. (B) Normalized expression of 8L:16D-induced transcripts (rDEI_8L:16D/16L:8D_ > 2) grouped by k-means clustering (see Methods). 16L:8D (red) and 8L:16D (blue) expression rhythms were transformed to Z-score together for clustering to retain relative magnitude. Black lines indicate median expression level. Grey rectangles indicate the dark portion of each photoperiod. The number of clusters is determined by the elbow method. Top enriched Gene Ontology (GO) and Kyoto Encyclopedia of Genes and Genomes (KEGG) pathways. (C) Hierarchical clustering of all 22810 transcripts by 16L:8D and 8L:16D patterns (Table S4). 16L:8D and 8L:16D expressions were transformed to Z score separately prior to clustering to obtain the pattern. Dendrogram edges are colored by the average rDEI_8L:16D/16L:8D_ of transcripts within the corresponding node. 132 clusters were defined by dynamic tree cutting. Clusters are indicated by the color bar: light grey and dark grey indicate clusters, and white indicates transcripts that are not assigned to any cluster. (D) Identification of photoperiod-induced clusters (average rDEI_8L:16D/16L:8D_ > 1.15 or average rDEI_8L:16D/16L:8D_ < 0.87). Statistical cutoff is drawn at adjusted *p*-value < 1.0 × 10^−20^ (one-sample Wilcoxon test; Bonferroni correction). Blue: 8L:16D-induced clusters; grey, other clusters. (E) Doughnut chart showing the overlap between clusters of 8L:16D-induced transcripts (rDEI_8L:16D/16L:8D_ > 2) and clusters of transcripts showing an 8L:16D-induction correlated pattern. (F) Expression pattern of transcripts in clusters 21, 25, and 26 normalized in each of the 16L:8D and 8L:16D dataset. Red: 16L:8D expression; blue; 8L:16D expression. Black lines indicate median expression level. Grey rectangles indicate dark period of each photoperiod (Table S4).

We next performed enrichment tests of Gene Ontology (GO) terms and Kyoto Encyclopedia of Genes and Genomes (KEGG) pathways from our 8L:16D-induced transcripts to understand the cellular pathways induced in winter photoperiods (Figure 1B and Table S2, S3) (Hvidsten, et al., 2001; Kanehisa and Goto, 2000; Ogata, et al., 1998). Supporting the validity of our approach, “photoperiod” and “red/far red light signaling” are enriched GO terms in the 8L:16D-induced transcripts from clusters A_W_-C_W_. Furthermore, the “response to carbohydrate,” “response to sucrose,” and “autophagy” GO terms and the “valine, leucine and isoleucine degradation” KEGG pathway are also enriched, highlighting that 8L:16D photoperiods signal the induction of energy response and nutrient conservation and scavenging pathways (Figure 1B and Table S2, S3). We also searched the list of putative winter transcripts for examples of photoperiod-specific function for the genes. *HOMOGENTISATE 1,2-DIOXYGENASE* (*HGO*-AT5G54080) from cluster A_W_ is an enzyme involved in tyrosine catabolism, specifically in winter photoperiods (Zhi, et al., 2016; Han, et al., 2013), and *MALATE SYNTHASE* (*MLS*-AT5G03860) from cluster C_W_ is a gene that is necessary for establishing true leaves in short winter-like days (Cornah, et al., 2004). Perhaps the clearest example of a gene that is important for winter development in the list is *TEMPRANILLO1* (AT1G25560), a transcriptional regulator that blocks flowering in winter-like photoperiods by repressing *FT* expression directly, in competition with CO (Johansson and Staiger, 2014; Castillejo and Pelaz, 2008). *TEMPRANILLO1* was also shown to have the hallmark expression pattern seen in the 8L:16D-induced cluster, A_W_ (Figure 1B).

### Defining an expression pattern for transcripts induced in short winter-like days

To determine whether the dark-phased expression pattern of winter photoperiod induced genes is linked to a high rDEI_8L:16D/16L:8D_, we performed hierarchical clustering of the normalized expression patterns for all transcripts from the 8L:16D and 16L:8D microarray experiments (Figure 1C and Table S4). This resulted in the identification of 131 expression pattern clusters. Three large clusters, numbered 21, 25, and 26 had expression patterns that appeared to be similar to clusters A_W_, B_W_, and C_W_ (Figure 1B) and also have statistically higher rDEI_8L:16D/16L:8D_ when compared to all of the transcripts represented by the microarray (Figure 1D). In particular, >85% of transcripts from cluster A_W_ fall within cluster 26, a large cluster of >1800 transcripts (Figure 1E-F). This congruence suggests that the temporal expression pattern represented by cluster 26 is correlated to higher rDEI_8L:16D/16L:8D_. We performed GO and KEGG analyses on clusters 21, 25, and 26 (Figure 1F). Cluster 26 contains terms that are similar to those found in clusters A_W_ and B_W_, including “photoperiodism”, “response to fructose”, and “vesicle-mediated transport” (a broader term containing “autophagy”). Cluster 26 also included the GO term “ubiquitin-like protein transferase activity” suggesting that the ubiquitin proteasome system is being induced in winter photoperiods and supporting the idea that cellular recycling programs are important winter processes.

### A winter gene, *PP2-A13,* is essential for Arabidopsis fitness in winter-like photoperiods

We previously curated a large group of genetic resources for F-box-type E3 ubiquitin ligases (Feke, et al., 2020; Feke, et al., 2019; Lee, et al., 2018; Lee C-M, 2017), which are part of the “ubiquitin-like protein transferase activity” GO term (Figure 1F). One of the winter F-box genes, *PP2-A13,* shares sequence similarity with the human lectin-containing F-box gene *F-BOX ONLY 2* (*FBXO2*, also known as *Fbs1/Nfb42/Fbx2/Fbg1*) which is critical for cytoplasmic glycoprotein quality control processes and results in age-related protein aggregation diseases when mutated in humans (Yoshida, et al., 2005; Dinant, et al., 2003; Yoshida, et al., 2003). The microarray data indicate that *PP2-A13* follows a dark-phased expression pattern similar to cluster A_W_ which we confirmed by qRT-PCR (Figure 2A). *PP2-A13* expression is qualitatively different between the 8L:16D and 16L:8D growth regimes. In both photoperiods there is a peak of expression near dawn that is subsequently repressed by exposure to light. In 8L:16D a second winter photoperiod-specific expression peak appears and is phased at about 4 hours after dusk.

**Figure 2.**
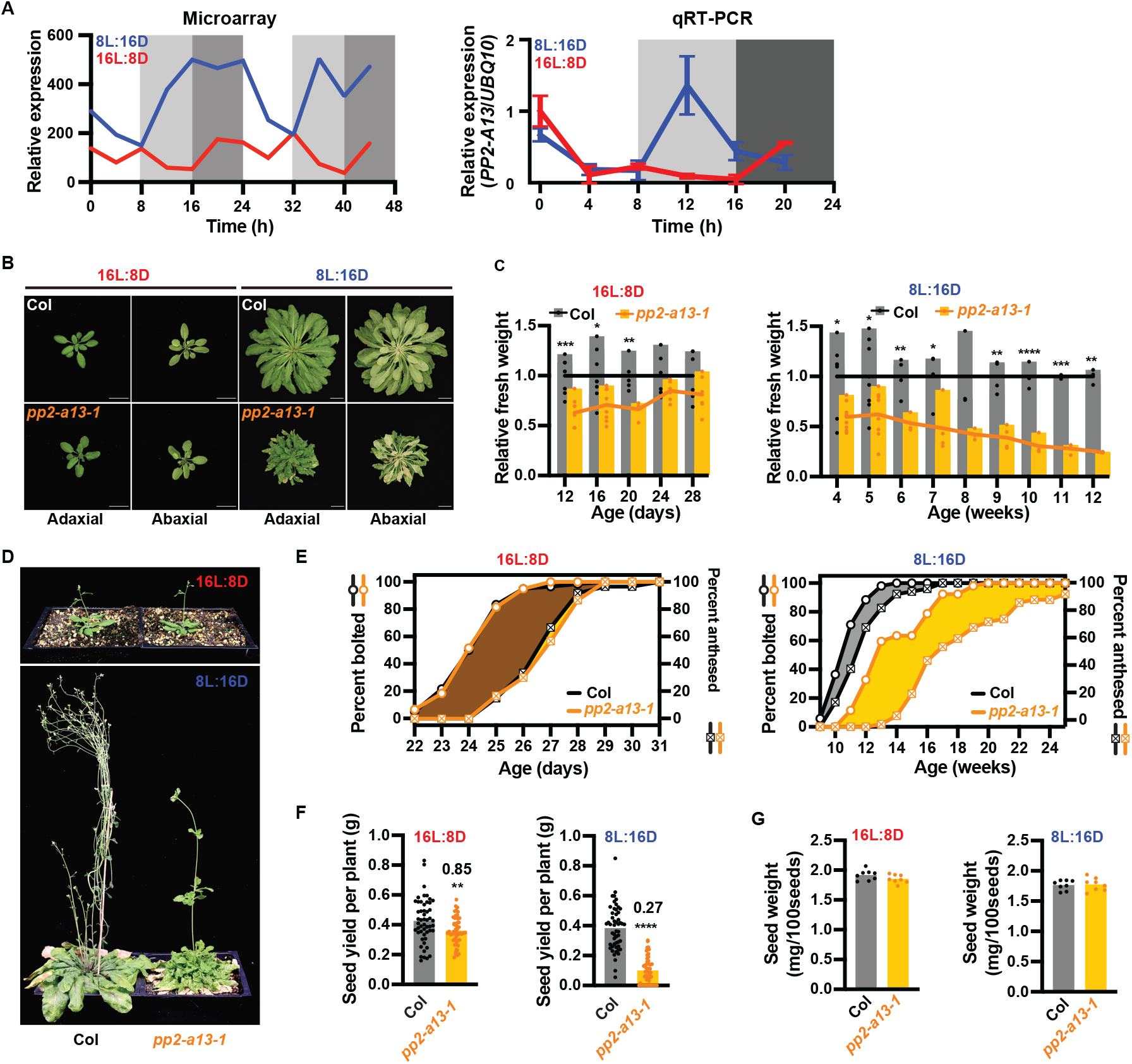
Disruption of the *PP2-A13* gene causes winter photoperiod-specific fitness defects. (A) Microarray expression data and qRT-PCR of *PP2-A13* from 12-day-old plants grown in 8L:16D (blue) and 16L:8D (red). n = 3 samples containing multiple seedlings for each time point. (B) Representative wild type (Col) and *pp2-a13-1* mutant plants. Grown for 24 days in 16L:8D or 11 weeks in 8L:16D. Adaxial and abaxial views of the rosettes are presented. Scale bar = 3 cm. (C) Aerial fresh weight of wild-type (Col) and *pp2-a13-1* mutant plants grown in 16L:8D and 8L:16D were normalized to the mean of wild-type (Col) at each time point. Black (Col wild type) or orange (*pp2-a13-1* mutant) lines indicate the mean of each genotype at different time points. *n* = 3-8 individual plants. Asterisks indicate significant difference between wild-type (Col) and *pp2-a13-1* mutant plants at each time point. *, *p*≤0.05; **, *p*≤0.01; ***, *p* β0.001; ****, *p*≤0.0001 (Welch’s t-test). (D) Representative wild-type (Col) and *pp2-a13-1* mutant plants grown for 28 days in 16L:8D or 14 weeks in 8L:16D. (E) Percentage of wild-type (Col) and *pp2-a13-1* mutant plants that are bolting or anthesed. Plants grown in 16L:8D (left) and 8L:16D (right). n = 52-60. (F) Total seed yield from wild-type (Col) and *pp2-a13-1* mutant plants grown in 8L:16D and 16L:8D. n = 52-60. (G) Seed weight in milligrams/100 seeds from wild-type (Col) and *pp2-a13-1* mutant plants grown in 8L:16D and 16L:8D. n = 8.

We next identified a transgenic line (*pp2-a13-1*) containing a T-DNA insertion in *PP2-A13* that has compromised expression of the *PP2-A13* transcript (Figure S2A-B). We assessed development over the life of the *pp2-a13-1* mutant in 8L:16D and 16L:8D growth conditions (Figure 2B-G and S2C-F). Strikingly, the leaves of the *pp2-a13-1* mutant senesce prior to flowering exclusively in 8L:16D, a qualitative reversal of these two important developmental processes (Figure 2B). In 8L:16D, the *pp2-a13-1* mutant is unable to maintain generation of biomass prior to flowering, while in 16L:8D the mutant is only partially compromised in biomass generation early in vegetative development and recovers later in development (Figure 2C and S2C-D). The phenotype of the mutant was complemented by expression of the full length *PP2-A13* driven by the native promoter confirming that the insertion in *PP2-A13* is causing the observed phenotypes (Figure S2G).

We then noted altered inflorescence morphology, bolting time, and anthesis in the *pp2-a13-1* mutant exclusively in 8L:16D (Figure 2D-E and S2E-F). Furthermore, in 8L:16D, 4 out of 52 (7.7%) mutant plants never underwent anthesis and did not produce seeds, while an additional 9 mutant plants produced no viable seeds (17.3%). We also found that the mutant plants in 16L:8D had a slight defect in seed yield while the 8L:16D grown mutant seeds were severely compromised, but neither growth condition caused a differential effect on weight per 100 seeds (Figure 2F-G). These results show that *PP2-A13* is necessary for Arabidopsis cellular health and reproduction in winter-like photoperiods.

### PP2-A13 works in parallel to autophagy and controls glycoprotein abundance

The cellular function of *PP2-A13* has not been studied in detail previously. We first determined the spatial pattern of expression of *PP2-A13* using a transgenic line expressing β*-glucuronidase* under the *PP2-A13* promoter (*PP2-A13*_*promoter*_::*GUS*) (Figure 3A). *PP2-A13* is expressed widely and does not seem to be tissue-specific. We then determined the subcellular localization of the PP2-A13 protein using transient expression of PP2-A13 fused to GFP in Arabidopsis protoplasts (Figure 3B). The PP2-A13 protein shows diffuse localization in the nucleus but also forms foci outside of the nucleus.

**Figure 3.**
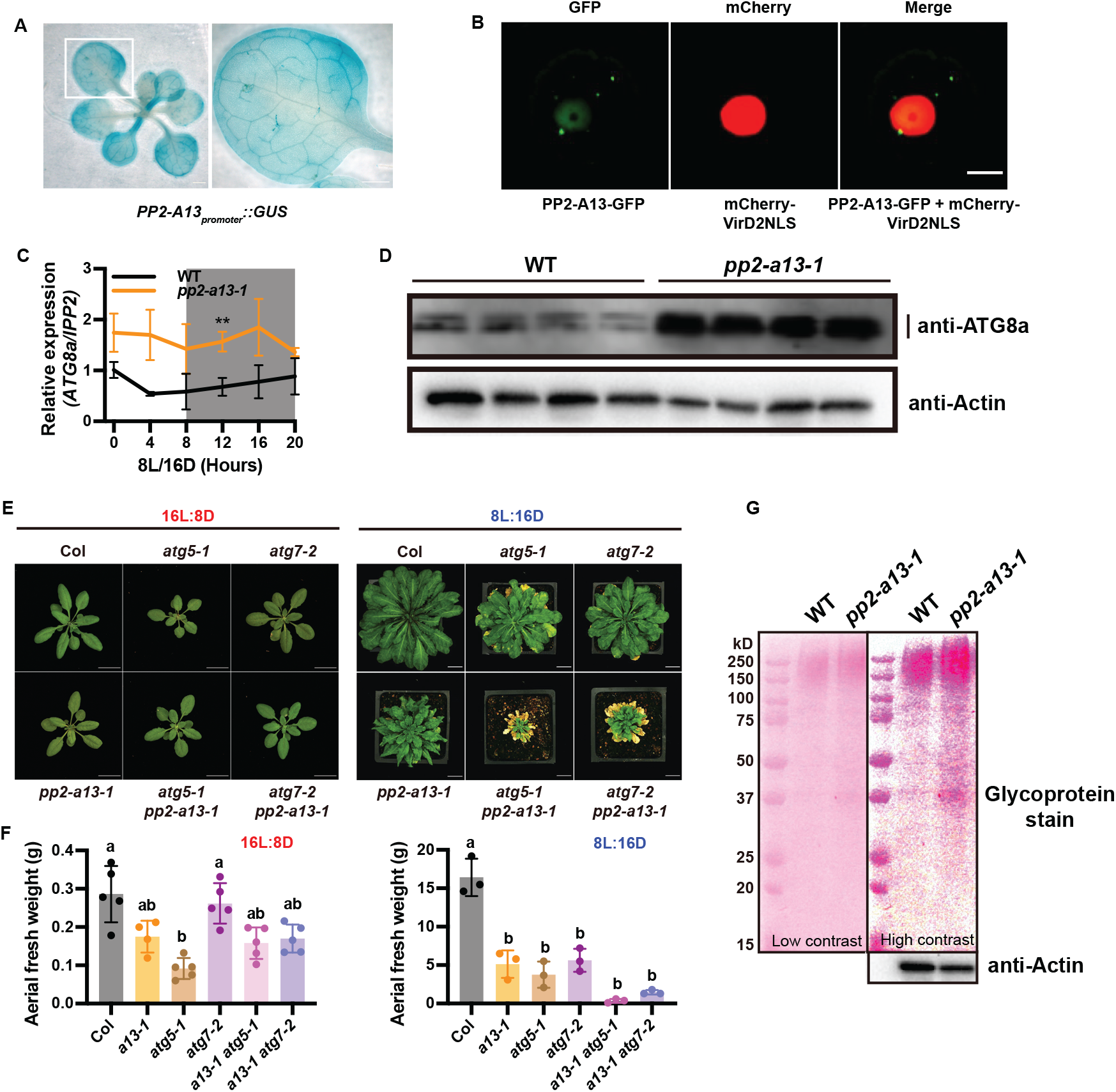
PP2-A13 works in parallel to autophagy and controls glycoprotein abundance. (A) GUS staining of the *PP2-A13*_*promoter*_::*GUS* transgenic line. The right image is the zoom-in view of the white box area in the left image. Scale bars = 1mm. (B) Subcellular localization of PP2-A13 was performed in Arabidopsis protoplasts. PP2-A13-GFP was co-expressed with a nuclear marker mCherry-VirD2NLS. Scale bar indicates 10 *μ*m. (C) qRT-PCR *ATG8a* from 6-week-old WT (black) and *pp2-a13-1* mutant (orange) grown in 8L:16D (blue). n = 3 individual samples for each time point. **, *p*≤0.01 (Welch’s t-test) (D) Immunoblot analysis of the *pp2-a13-1* mutant. Crude protein extracts of 11-week-old wide-type (Col) and *pp2-a13-1* mutant were subjected to SDS-PAGE and immunoblot analysis with anti-ATG8a antibody. Equal protein loads were confirmed by immunoblot analysis with anti-Actin antibody. (E) Representative images of wild-type (Col), *pp2-a13-1*, *atg5-1*, *atg7-2*, *atg5-1 pp2-a13-1*, and *atg7-2 pp2-a13-1* mutant plants grown in 16L:8D for 28 days or 8L:16D for 87 days. Scale bar = 2 cm in 16L:8D and 3 cm in 8L:16D. (F) Aerial fresh weight of wild-type (Col), *pp2-a13-1*, *atg5-1*, *atg7-2*, *atg5-1 pp2-a13-1*, and *atg7-2 pp2-a13-1* mutant plants grown in 16L:8D and 8L:16D. Different letters indicate statistically significant differences as determined by one-way ANOVA followed by Dunnett’s T3 multiple comparison test; *p*≤0.05. Error bars indicate SD (n = 3-5 individual samples). (G) Glycoprotein analysis for WT and *pp2-a13-1* mutant. Crude protein extracts of 11-week-old wide-type (Col) and *pp2-a13-1* mutant were subjected to SDS-PAGE and stained with Pierce Glycoprotein Staining Kit. Equal protein loads were confirmed by immunoblot analysis with anti-Actin antibody.

The phenotypic effects of the *pp2-a13-1* mutant are reminiscent of the effects of autophagy mutants grown in short winter-like photoperiods. The “autophagy” GO term is enriched in our winter gene list, and autophagy is critical for nutrient recycling and cellular health in short days in Arabidopsis (Izumi, et al., 2013). It is possible that *PP2-A13* participates in autophagy by mediating ubiquitylation of targets for selective autophagy. Indeed, in the *pp2-a13-1* mutant plants the expression of *ATG8a* mRNA is induced and the ATG8a protein is more highly accumulated, similar to the effects seen in autophagy mutants (Figures 3C-D) (Phillips, et al., 2008). To test if the *pp2-a13-1* phenotypes are due to defects in autophagy, we crossed the *pp2-a13-1* mutant with the *atg5-1* and *atg7-2* mutants and observed the phenotypes of the double mutants (Figures 3E-F). The double mutants showed defects in growth that were more severe than either single mutant alone, exclusively in short winter-like days. This indicates that PP2-A13 functions in a pathway that is parallel to autophagy.

Based on work done with lectin-containing F-box proteins in mammalian systems, we hypothesized PP2-A13 may function to control glycoprotein abundance (Yoshida, et al., 2019). We tested this by examining the levels of glycosylated proteins in the *pp2-a13-1* mutant in plants grown in short winter-like photoperiods (Figure 3G). We found that the abundance of glycosylated proteins was higher in the mutant plants suggesting a conservation of function with mammalian lectin-containing f-box proteins.

### *PP2-A13* expression is photoperiodically induced

Due to the importance of *PP2-A13* in plant winter survival, we wanted to create a reporter system to rapidly explore the underlying systems that controls winter-photoperiod expression of *PP2-A13*. To achieve this, we generated transgenic plants expressing the *Luciferase* gene under the control of the *PP2-A13* promoter (*PP2-A13*_*promoter*_::*luciferase*) (Figure 4A). We measured luminescence from the *PP2-A13*_*promoter*_::*luciferase* plants under 8L:16D and 16L:8D conditions (Figure 4B). The patterns generated from this experiment were similar to those seen in the qRT-PCR and microarray experiments (Figure 2A). In 8L:16D, the reporter line shows the winter-photoperiod specific expression peak after dusk, while in both 8L:16D and 16L:8D the reporter line shows the dawn expression peak and subsequent repression by light exposure. To examine the daily expression shape and compare across experiments, we normalized the data to the trough and peak levels. While this removes amplitude information, it gives a clearer view of the comparative expression pattern shapes (Figure 4C). We also calculated the rate of change in intensity (“intensity change”) (Figure 4D). These analyses confirm the winter specific expression peak of *PP2-A13* and show that *PP2-A13* expression rises rapidly after dusk in 8L:16D and slowly in 16L:8D.

**Figure 4.**
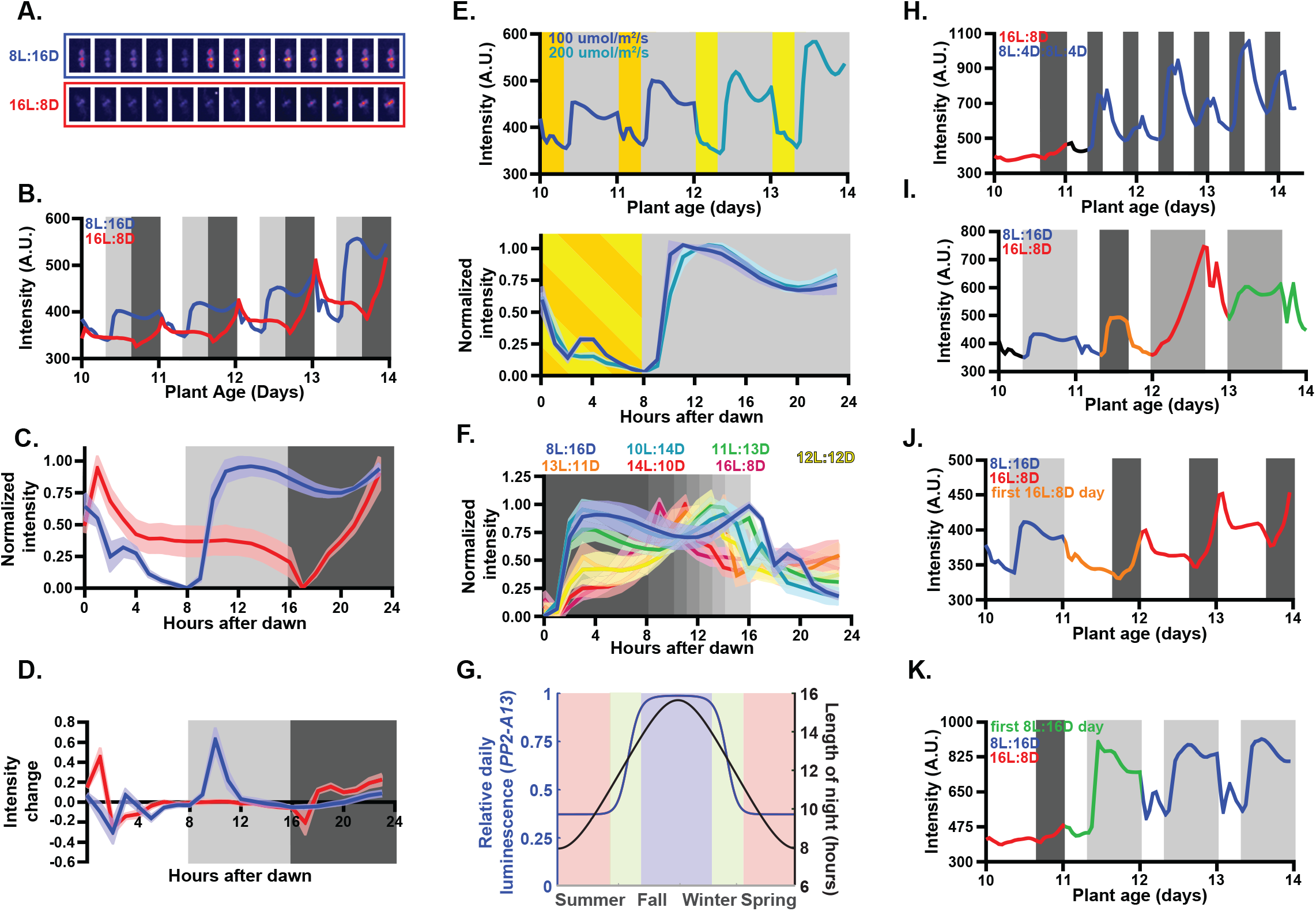
A winter photoperiod measuring mechanism controls winter gene expression. (A-C) *PP2-A13*_*promoter*_::*Luciferase* expression from plants grown under 8L:16D and 16L:8D photoperiods. Grey shading represents the dark period for the various photoperiod experiments. Colored lines represent the intensity traces and shading represents the standard deviation. Black traces in the raw luciferase intensity plots represent time periods that were excluded from normalization and rate calculations. (A) False color images of representative plants taken every two hours from ZT0 to ZT24. (B) Average from traces of raw luciferase intensity. (C) Normalized traces of the daily luciferase intensity pattern. (D) Average rate of change in expression. (E) Plants grown under 8L:16D with 100 μM m^−2^ s^−1^ light (dark yellow) conditions were transferred into 200 μM m^−2^ s^−1^ light (light yellow). Note that for this experiment, the false-colored 100 μM m^−2^ s^−1^ ZT24 image is the same as the 200 μM m^−2^ s^−1^ ZT0 image. (F) Determination of the critical photoperiod. Traces are from plants grown in indicated conditions. The individual traces are presented in figure S4A and rates are presented in figure S3C. (G) Night lengths in Landsberg, Germany (black) and estimated yearly expression pattern (red) of *PP2-A13*_*promoter*_::*Luciferase* as calculated from the normalized expression in figure 3F. (H) Plants grown under 16L:8D conditions were transferred to double dusk (8L:4D:8L:4D) conditions on day 11. Individual movie frames, normalized pattern, and average rates are presented in figure S3D. (I) Plants were grown under 8L:16D conditions until day 10. On day 11, plants underwent a dawn phase advance of 8 hours but kept in 8L:16D for the remainder of the experiment. Normalized plots were excluded from this figure but rates are presented in figure S3E. (J) Plants grown under 8L:16D conditions were transferred into 16L:8D conditions on day 11. Individual movie frames, normalized pattern, and average rates are presented in figure S3F. Note that for this experiment, the false colored 8L:16D ZT24 picture is the same as the 16L:8D ZT0 picture. (K) Plants grown under 16L:8D conditions were transferred into 8L:16D conditions on day 11. Individual movie frames, normalized pattern, and average rates are presented in figure S3G. Note that for this experiment, the false colored 16L:8D ZT24 picture is the same as the false colored 8L:16D ZT0 picture.

We next tested whether *PP2-A13* expression is under the control of a “true” photoperiodic measuring system independent of light intensity. We grew the plants in 8L:16D at 100 μM m^−2^ s^−1^ (8L_100_:16D) for the first part of the experiment and then on day 12 we maintained day length but doubled the light intensity to 200 μM m^−2^ s^−1^ (8L_200_:16D) (Figure 4E and S3A), matching the daily light integral of the 16L:8D experiment in figure 4B. The pattern of *PP2-A13* expression was nearly unchanged after doubling the light intensity. We also performed the entire experiment with plants grown in 8L_100_:16D and 8L_200_:16D and did not detect a difference in the pattern of *PP2-A13* expression (Figure S3B). This indicates that the expression pattern of *PP2-A13* is reporting on a true photoperiod measuring mechanism that operates independent of light intensity.

To determine the critical photoperiod in which *PP2-A13* expression changes from the winter-like pattern to the summer-like pattern, we imaged the reporter plants in photoperiods ranging from 4L:20D to 20L:4D (Figure 4F, S3C, and S4A). Plants grown in photoperiods with longer nights, akin to fall and winter (8L:16D, 10L:14D, and 11L:13D), exhibit the hallmark *PP2-A13* winter expression signature. Plants grown in photoperiods with days at least one hour longer than night, akin to late spring and early summer (14L:10D, 16L:8D), exhibit summer photoperiod-like expression patterns. These trends continue in more extreme photoperiods (4L:20D and 20L:4D) as well (Figure S4B-C). In plants grown in photoperiods with days that are equal to or slightly longer than nights, akin to spring or fall equinox and early spring or late summer (12L:12D, 13L:11D), the expression pattern appears to be in a transitional state with a small expression “shoulder” early in the night, suggesting that these are near the critical photoperiod.

We next wanted to know how this expression pattern may translate to levels of *PP2-A13* across one year. We calculated the area under the curve for each experiment from the critical photoperiod data (Figure S4A) and fit this data to a curve with an approximate sigmoid function (Figure S5). We then determined the night lengths over one year in central Germany, where the Columbia ecotype was first isolated (Latitude 48° N), and used this information to calculate a predicted expression level for *PP2-A13* over one full year (Figure 4G). The data clearly shows the expression pattern of *PP2-A13* is not linear with the night length, clearly demonstrating a photoperiodic switch in expression levels.

Using the real-time reporter we can observe post-dusk induction rates before and after the critical photoperiod in the same 24 hour period (a “double dusk” experiment), a direct test of Bünning’s two state model. We performed this experiment by growing the reporter plants in 16L:8D and then exchanging the light cycle with 8L:4D:8L:4D, maintaining the same daily light integral as 16L:8D but providing one dusk prior to the critical photoperiod and one after the critical photoperiod (Figure 4H and S3D). Supporting a two state model, the rate of induction and expression peak are higher in the first dark period than the second dark period. This, along with the critical photoperiod study (Figure 4F), shows that the plant is transitioning between two dark response states across the 24-hour day.

Circadian clock or hourglass-like timers function in photoperiodic measurement systems (Bradshaw and Holzapfel, 2010; Saunders, 2005; Saunders, 1997). A circadian clock-like mechanism takes time to re-entrain to a new dawn after a phase shift while an hourglass, by nature, resets immediately to a new dawn. We grew the plants in 8L:16D and then advanced the phase of dawn by eight hours. Subsequent to the phase advance, we maintained the 8L:16D photoperiod (Figure 4I and S3E). On day one after the phase advance (Figure 4I, red trace), we observe a *PP2-A13* expression pattern that is different than any daily expression pattern observed in previous experiments. On day two after the phase shift (Figure 4I, green trace) the expression pattern is similar to the standard 8L:16D pattern seen previously. This suggests that the two dark response states controlling photoperiodic *PP2-A13* expression are under the control of a circadian clock-like timer.

Photoperiodic timing mechanisms often count the number of hours of dark or the number of hours of light rather than the relative day and night lengths (Lumsden and Millar, 1998; Vince-Prue, 1975). To determine if winter gene expression is measuring the length of day or length of night, we performed photoperiod shift experiments. We grew plants in 8L:16D and then changed the light cycle to 16L:8D and vice versa (Figure 4J-K and S3F-G). In both experiments, on the first day after the shift the expression patterns reset to the new photoperiod. The plants are able to readjust the post-dusk expression pattern after only experiencing one light period, suggesting that this process counts the number of hours of light.

*CONSTANS* (*CO*) mediates the photoperiodic induction of some genes in long days in Arabidopsis, including the florigen *FT*. Our results show that the winter photoperiod transcript induction system is phased to the early part of the 24 hour day which is opposite to *CO*. We tested whether the CO photoperiod measuring system controls winter transcript induction. We crossed the *co-9* mutant into our reporter and grew the plants in 16L:8D and 8L:16D for imaging (Figure 5A-B). The expression pattern of the reporter was nearly identical in the wild-type and *co-9* mutant plants despite the *co-9* mutant plants flowering later than the wild-type plants. This strongly indicates that the photoperiod measuring mechanism is distinct from the mechanism that controls photoperiodic flowering. In support of this idea, our *PP2-A13_promoter_::GUS* transgenic line does not show vein specific expression, the tissue where the CO/FT mechanism functions (Figure 3A) (An, et al., 2004).

**Figure 5.**
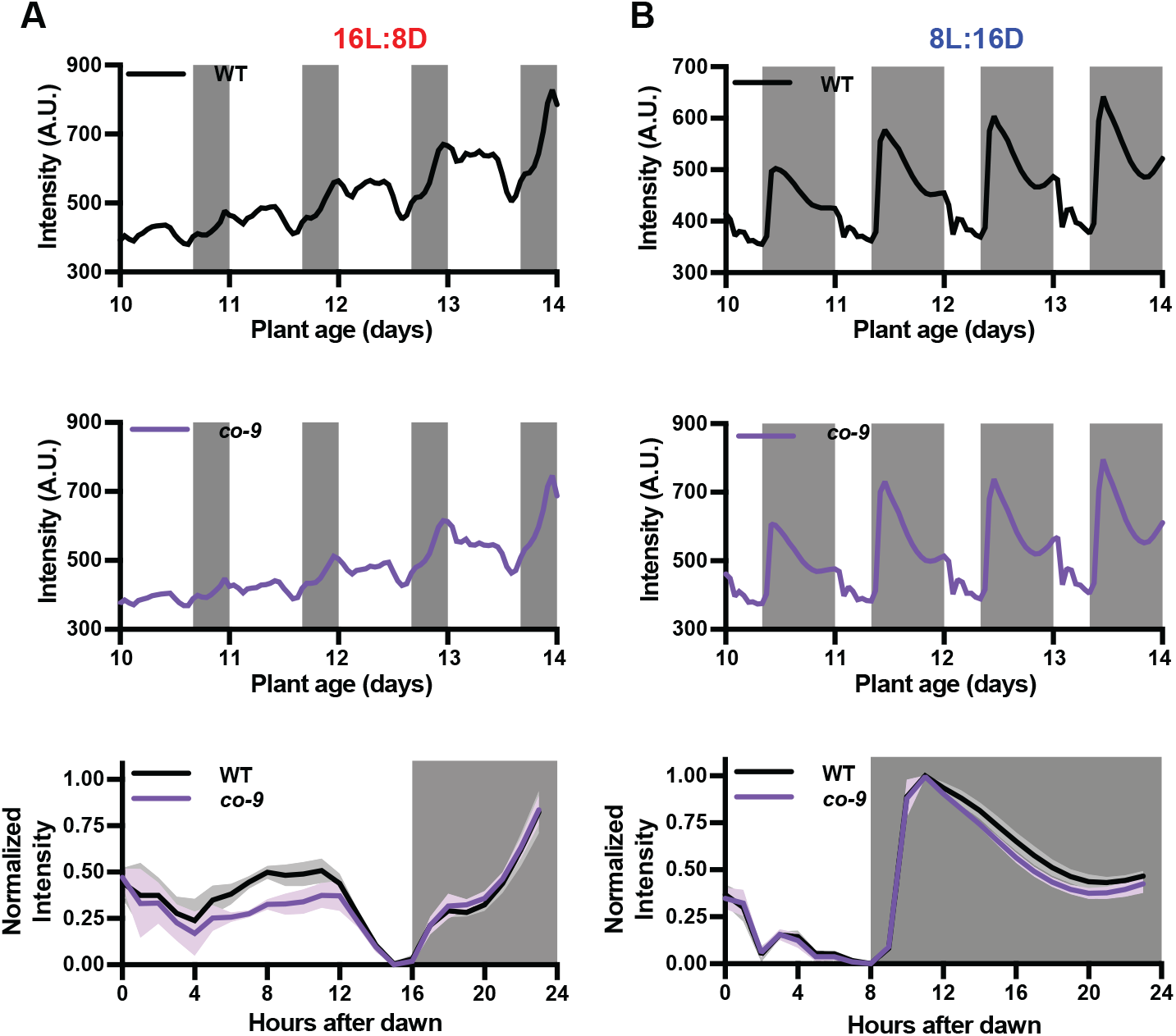
CONSTANS does not regulate the photoperiodic induction or repression of winter genes. (A-B) *PP2-A13*_*promoter*_::*Luciferase* traces and normalized traces from wild-type and *co-9* mutant plants grown under (A) 16L:8D and (B) 8L:16D photoperiods.

### Darkness is transmitted through the photosynthetic apparatus to photoperiodic induction of winter genes

A necessary component of a photoperiod measuring mechanism is a sensor(s) that can distinguish between light and dark. Plants sense light/dark transitions through the photosynthetic apparatus or environmental sensing photoreceptors. To determine whether photosynthesis or photoreceptors are sensing light/dark transition to control *PP2-A13* expression, we replaced the first eight hours of darkness in an 8L:16D growth condition with red light (635 nm), a single photosynthetically active wavelength that is sensed by phytochromes, red-light photoreceptors, in plants. This regime was performed at two red light intensities, one at 100 μM m^−2^ s^−1^ in which phytochrome signaling is presumably saturated and the intensity is well above the light compensation point (8L:8R_100_:8D), and the second at 5 μM m^−2^ s^−1^ in which phytochrome signaling should be active but is well below the light compensation point for Arabidopsis (the 8L:8R_5_:8D) (Figure 6A and S6A) (Moraes, et al., 2019). In the 8L:8R100:8D condition, *PP2-A13* expression remains low when the lights change to red, similar to the pattern seen in 16L:8D and showing that high red light is sufficient to mimic white light in control of *PP2-A13* expression. However, in the 8L:8R_5_:8D condition, the expression pattern is similar to the 8L:16D winter photoperiod expression pattern. This shows that light is sensed by a system that requires light intensity above the compensation point.

**Figure 6.**
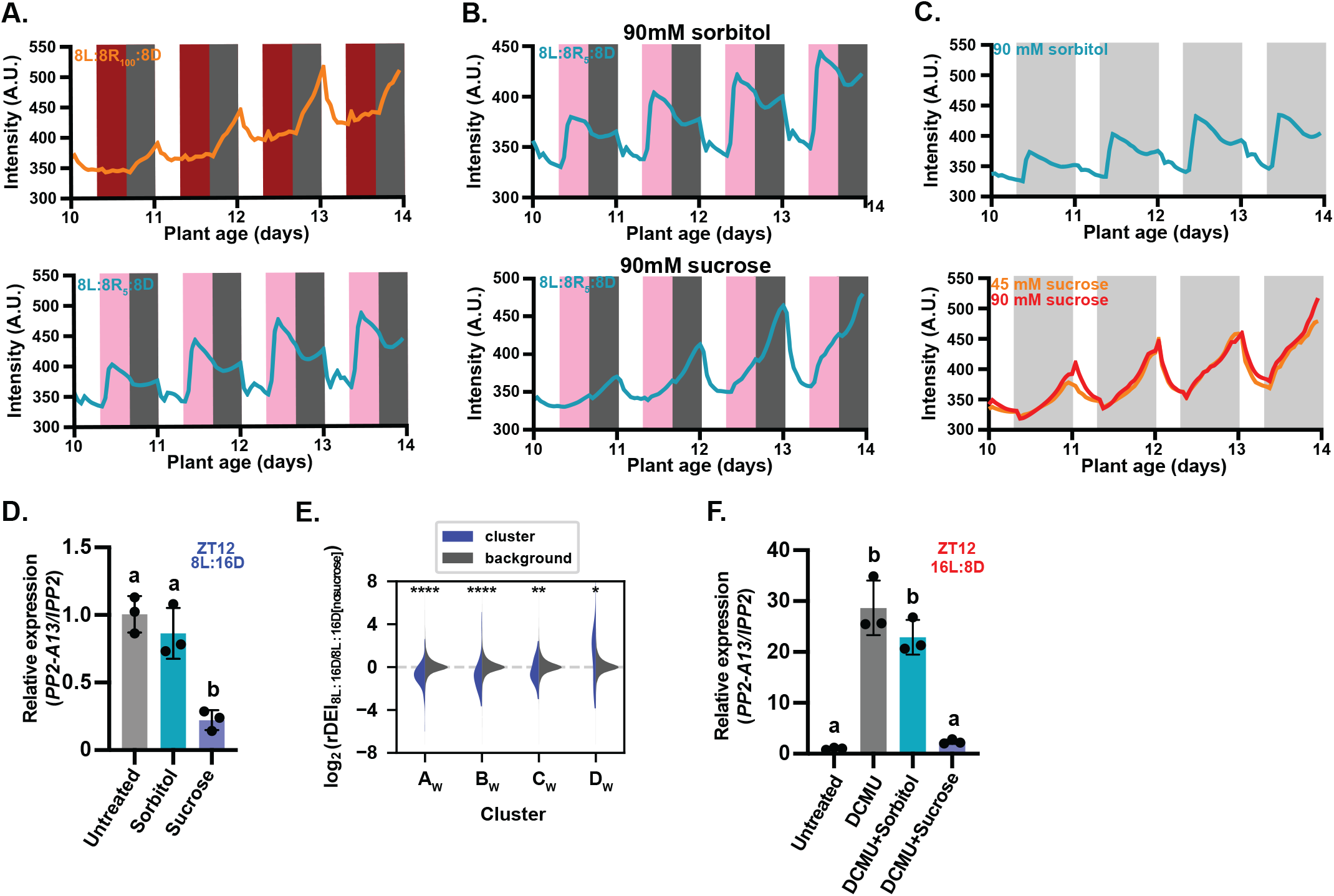
The photosynthetic apparatus senses darkness for winter photoperiod time measurement. (A-C) *PP2-A13*_*promoter*_::*Luciferase* trace data from plants grown in (A) 8L:8R100:8D (top panel) and 8L:8R5:8D (bottom panel), (B) 8L:8R5:8D treated with 90mM sorbitol (top panel) and 8L:8R5:8D treated with 90mM sucrose (bottom panel), (C) 8L:16D treated with 90mM sorbitol (top panel) and 8L:16D treated with 90mM sucrose (bottom panel). (D) qRT-PCR of *PP2-A13* from 12-day-old plants grown in 8L:16D. The indicated treatment started at ZT0 and tissue was collected at ZT12. Means with different letters are significantly different determined by one-way ANOVA followed by Dunnett’s T3 multiple comparison test; *p*≤0.05. Error bars indicate SD (n = 3 samples containing multiple seedlings). (E) The rDEI_8L:16D sucrose/8L:16D_ no sucrose of 8L:16D-induced transcripts (blue) in comparison to the rDEI_8L:16D sucrose/8L:16D_ no sucrose of all other transcripts (grey). rDEI_8L:16D sucrose/8L:16D_ no sucrose is calculated as the rDEI of the DIURNAL “shortday” time course divided by the rDEI of the DIURNAL “LER_SD” time course. Asterisks indicate statistical significance between the 8L:16D-induced cluster and the background: *, *p*<0.05; **; *p*<0.01; ***, *p*<0.0005; ****, *p*<0.0001 (Welch’s t-test; Bonferroni correction). (F) qRT-PCR of *PP2-A13* from 12-day-old plants grown in 16L:8D. The indicated treatment started at ZT0 and tissue was collected at ZT12. Different letters indicate statistically significant differences as determined by one-way ANOVA followed by Dunnett’s T3 multiple comparison test; *p*≤0.05. Error bars indicate SD (n = 3 samples containing multiple seedlings).

One of the main products of photosynthesis in Arabidopsis is sucrose. To test if sucrose can alter the *PP2-A13* photoperiodic response, we performed imaging experiments in 8L:8R_5_:8D in the presence of exogenously supplied sucrose (Figure 6B and S6B). The winter photoperiod expression peak of *PP2-A13* is nearly ablated when sucrose is supplied to the plants and begins to resemble the expression pattern seen in summer photoperiods. We also tested this in white light with two concentrations of sucrose, both of which suppressed the winter expression peak (Figure 6C and S6C). We then tested the repression of the winter expression peak using qRT-PCR. We grew plants in 8L:16D and treated them with sucrose or sorbitol starting at ZT0. We collected tissue at ZT12 (4 hours post-dusk in 8L:16D), and measured *PP2-A13* expression (Figure 6D). We found that the sorbitol treatment had little effect on *PP2-A13* expression while the sucrose repressed expression, similar to what we found with the reporter. These results show that sucrose, an important product of photosynthesis, can suppress the winter photoperiod expression of *PP2-A13*. Furthermore, the three night-phased clusters of winter genes, A_W_, B_W_, and C_W_ (Figure 1B), are all repressed by the presence of sucrose in the growth media (Figure 6E). This result supports the idea that winter transcripts are generally repressed by sucrose in 8L:16D growth conditions, as observed with *PP2-A13*.

Our results indicate that the photosynthetic apparatus senses darkness to control winter gene induction. To further test this idea, we grew plants in 16L:8D but blocked photosynthesis using a specific chemical inhibitor of photosystem II called 3-(3,4-dichlorophenyl)-1,1-dimethylurea (DCMU). It was technically challenging to perform this experiment using our real-time reporter, necessitating the use of qRT-PCR. In 16L:8D we treated the plants with DCMU at ZT0 of day 12 (Figure 6F). We then collected tissue at ZT12 when the plants should have very low expression of *PP2-A13* because they are still in the light. In the presence of DCMU, *PP2-A13* expression is induced, despite being the light. This effect was reversed upon the addition of sucrose. This result strongly indicates that darkness, with respect to *PP2-A13* expression, is being sensed by the inactivity of the photosynthetic apparatus rather than phytochrome, cryptochrome, or other photoreceptors.

### Rhythmic starch controls the phasing of the winter-photoperiod measuring mechanism

We next wanted to determine which process acts to differentially interpret the darkness across a 24-hour day. This mechanism sets the transition of the plant between “state1” and “state2” (Figure 4F and 4H). Extensive studies have shown that starch production and breakdown is circadian clock and photoperiod regulated and controls a large host of rhythmic metabolic processes in and out of the chloroplast (Kim, et al., 2017; Mengin, et al., 2017). Furthermore, starchless mutants in Arabidopsis, such as *phosphoglucomutase* (*pgm*) mutants, have more severe growth and developmental defects in winter photoperiods than in summer or equinox photoperiods (Eimert, et al., 1995). To test whether rhythmic starch production is controlling the state 1/state2 for *PP2-A13* expression, we crossed the *PP2-A13*_*promoter*_::*luciferase* reporter into the *pgm* mutant and monitored expression in 8L:16D, 16L:8D, and 8L:4D:8L:4D growth conditions (Figure 7A-C-pink traces). The *pgm* mutant causes altered expression of *PP2-A13* in all three conditions. In 8L:16D, the winter expression peak is delayed to near the middle of the night (Figure 7A). In 16L:8D in the *pgm* mutant, *PP2-A13* expression now is more rapidly induced and has two peaks of expression, similar to wild type in short winter-like days (Figure 7B). This was confirmed in a sucrose treatment experiment that shows the ablation of the first peak and restoration of the standard expression pattern seen in 16L:8D in wild type(Figure S7A-B). These results indicate that the underlying rhythmic process that defines state1 and state2 is misphased and delayed to a later part of the 24 hour day in the *pgm* mutant. The 8L:4D:8L:4D condition tests this more directly (Figure 7C). This experiment shows that the two-state system that exists in wild-type plants has been changed in the *pgm* mutant so that there is no distinction between darkness early or late in the 24-hour day, suggesting that the *pgm* mutant lacks the ability to accurately control winter gene expression.

**Figure 7.**
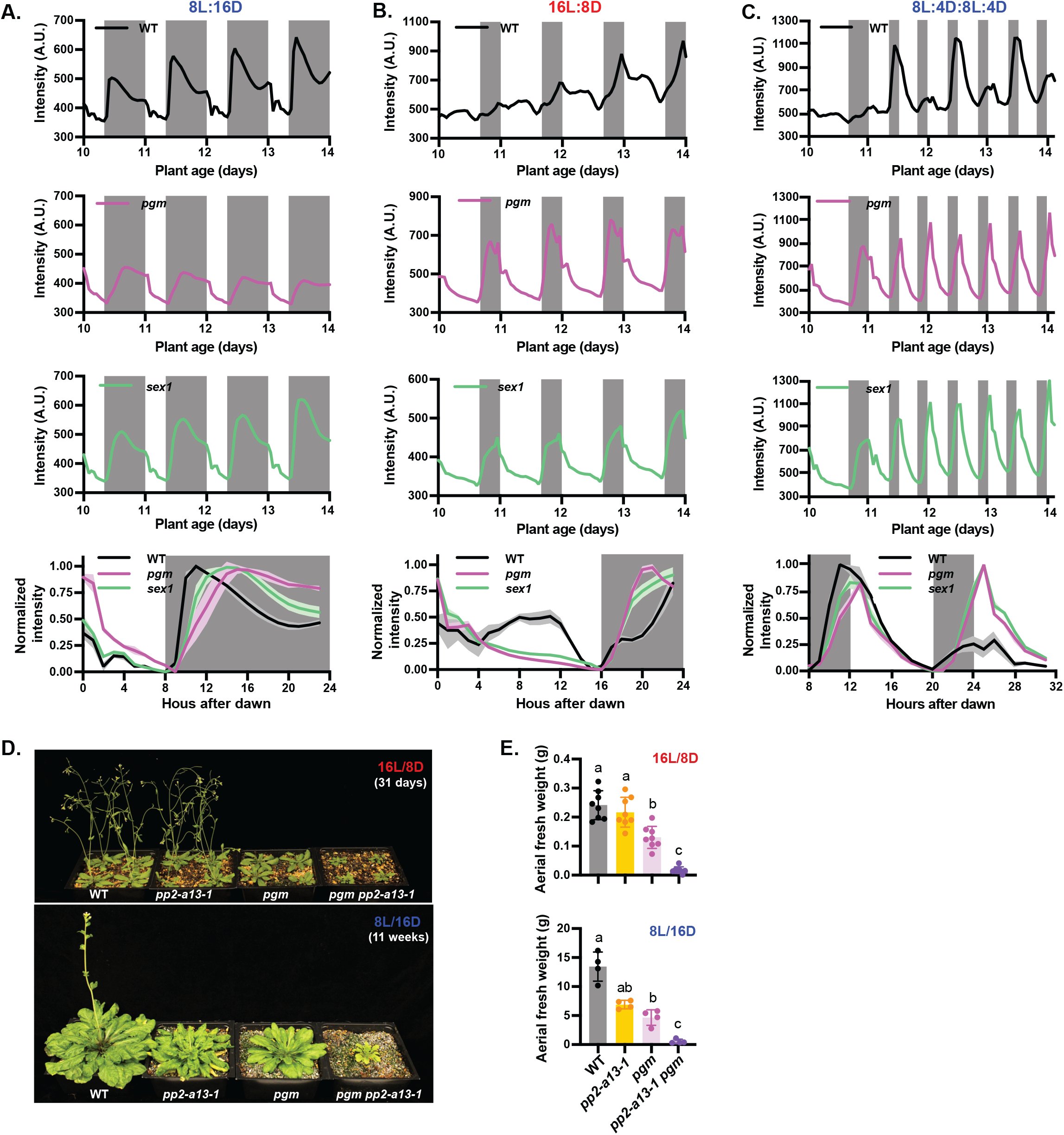
A metabolic coincidence mechanism controls winter photoperiod gene expression. (A-C) Average and normalized *PP2-A13*_*promoter*_::*Luciferase* traces from wild type, *pgm* mutant, and *sex1* mutant grown in (A) 8L:16D, (B) 16L:8D, and (C) 8L:4D:8L:4D. Note that for this experiment, the traces and average of WT in 8L:16D is the same as the WT in figure 5. (D) Representative wild-type (Col), *pp2-a13-1* mutant, *pgm* mutant, and *pgm pp2-a13-1* double mutant plants grown for 31 days in 16L:8D or 11 weeks in 8L:16D. (E) Aerial fresh weight of wild-type (Col), *pp2-a13-1* mutant, *pgm* mutant, and *pgm pp2-a13-1* double mutant plants grown for 25 days in 16L:8D or 11 weeks in 8L:16D. Different letters indicate statistically significant differences as determined by one-way ANOVA followed by Dunnett’s T3 multiple comparison test; *p*≤0.05. Error bars indicate SD (n = 4-8 individual plants).

The effects of *pgm* on winter gene expression could be explained by a lack of rhythmic starch production or alternately the low starch levels of the mutants. To test this we crossed the *PP2-A13* reporter into the *starch excess1* (*sex1*) mutant which maintains high levels of starch (Caspar, et al., 1991). We again monitored expression in 8L:16D, 16L:8D, and 8L:4D:8L:4D growth conditions and found a similar result as the *pgm* mutant (Figure 7A-C-green traces). *PP2-A13* expression is induced in 8L:16D but delayed when compared to wild type (Figure 7A). In 16L:8D *PP2-A13* is also induced rapidly and the induction can be suppressed by sucrose (Figure 7B and S7A-B). Again, state 1 and state 2 are altered in the 8L:4D:8L:4D growth condition showing that the plant can’t distinguish between winter and summer photoperiods (Figure 7C). This result suggests that starch levels are not being measured by the plant, but rather rhythmic starch production and breakdown maintains the phasing of a downstream rhythmic metabolic product, gene, protein, or other biological molecule that differentiates between dusk that occurs in state 1 and state 2.

We can further test this idea using the *pp2-a13-1* mutant. The previous result suggests that the *pgm* mutant is inappropriately activating winter genes, such as *PP2-A13*, in summer and winter photoperiods. Thus, the *pp2-a13-1* mutant phenotype would be apparent in the *pgm* mutant line in both summer and winter photoperiods, rather than exclusively in winter photoperiods like the wild-type plants. We crossed the *pgm* mutant with the *pp2-a13-1* mutant and found growth defects in both winter and summer photoperiods in the double mutant plants (Figure 7D-E). This is clearly seen in the representative images of the plants and is quantified in the fresh weight measurements. This result confirms the idea that rhythmic starch production is necessary for plants to measure seasons and that photosynthesis and rhythmic starch converge to form a metabolic coincidence mechanism to control winter gene expression.

## Discussion

Plants have been one of the preeminent study systems for understanding photoperiod measuring mechanisms for more than one hundred years (Lumsden and Millar, 1998; Vince-Prue, 1975; Bunning, 1969). This is because of the visually stunning transition from vegetative growth to flowering, which is often under tight control of a photoperiod measuring mechanism. Despite this, the intense focus on photoperiodic flowering has come at the cost of searching for additional photoperiod measuring mechanisms and understanding the full scope of biological processes that are enacted throughout the year. Winter can appear to be a time of inactivity for plants, but here we clearly show that plants are actively promoting the expression of genes to maintain fitness in winter photoperiods.

Here we describe the photoperiodic control of winter gene expression and show that it relies on a type of external coincidence we term “metabolic coincidence”. In this mechanism we show that darkness, sensed through the photosynthetic apparatus, is differentially interpreted by a process controlled by rhythmic metabolism downstream of starch production. This mechanism is distinct from, and functions opposite to, the CO/FT photoperiod measuring system for flowering in Arabidopsis. Interestingly, photosynthesis and starch metabolism both occur in the chloroplast of Arabidopsis, making it possible that this system resides in any chloroplast-containing cell in the plant, rather than being restricted to transport tissues like the CO/FT mechanism. This is supported by the expression pattern of the *PP2-A13*_*promoter*_::*luciferase* and *PP2-A13*_*promoter*_::*Gus* reporters, which do not show vein-specific expression.

Here we have identified the two main cellular systems that coordinate to form a seasonal measurement system, but in future work we will likely need to identify many more molecular players that participate in this process. It will be critical to identify whether photosynthetic redox signaling or lack of photosynthetic carbon capture is providing the dark signal that triggers rapid winter transcript activation (Foyer, 2018). It will also be important to identify the gene, protein, or molecule that is phased by rhythmic metabolism to enact gene expression in winter photoperiods. Furthermore, this system resides in the chloroplast but manifests as gene expression changes in the nucleus. We will need to determine how the signal is communicated between these two cellular compartments, especially the exact transcription factors that are involved. The real-time luciferase reporter, akin to the first real-time circadian clock reporters (Millar, et al., 1995a; Millar, et al., 1995b; Millar, et al., 1992), paves the way for identifying these components using a host of genetic and reverse genetic approaches

Winter transcripts in plants include many genes involved in cellular recycling, energy conservation, amino acid catabolism, growth cessation, and dormancy (Fig.1B and 1F). The plant is actively promoting mechanisms to protect itself from starvation in a low energy condition. Furthermore, the dark-response rhythm can be ablated by providing an exogenous energy source to the plant, and the winter photoperiod measurement mechanism relies on darkness being sensed by the photosynthetic apparatus. This indicates that there is an intimate connection between the energy state of the plant and its ability to enact this seasonal developmental program. Thus, it may be apropos in this case to refine the photophilic and skotophilic nomenclature that was proposed for photoperiodic flowering. In the case of winter transcripts it may be easier to imagine that when an early dusk occurs the plant is afraid to starve, and thus the plant is in a famophobic state. When dusk occurs late in the day the plant is afraid to inappropriately conserve and not spend its resources and thus is in a conservaphobic state.

We chose to focus our attention on the study of one winter gene, *PP2-A13*, because the insertion mutant line has striking and easily observable developmental defects (Figure 2 and S2). Here we show that *PP2-A13* functions in a plant cellular pathway that is parallel to autophagy and likely helps promote degradation of glycosylated proteins, akin to human lectin-containing F-box proteins. It will now be important to further define the sugar-binding specificity and scope of potential targets of *PP2-A13* to refine our understanding of its function expand our knowledge of the cellular pathways that it controls. It will also be important to further explore the relationship between PP2-A13 and autophagy to understand how they communicate, whether they share conserved targets, and understand their winter-photoperiod specific roles in plants.

Seasonal biological cycles of plant development are at the core of healthy ecosystems on earth, with plants acting as primary producers. Plants predict both adverse and beneficial seasonal changes by measuring photoperiod, but climate change is rapidly decoupling photoperiod from important seasonal cues such as temperature and water availability (Inoue, et al., 2020; Walker, et al., 2019; Stromme, et al., 2017; Fournier-Level, et al., 2016; Diez, et al., 2014). Importantly to our work, climate change has a disproportionately large effect on winter (Kreyling, 2010) and many plants need winter signals for proper reproductive and vegetative development. It is critical that we continue to explore the conservation of winter photoperiodic measurement mechanisms to ensure future robustness of our most important crops in the face of climate change.

## Supporting information

Supplementary Codes

## ACKNOWLEDGEMENTS

We would like to thank Christopher Adamchek and Suyuna Eng Ren for their technical support. We would also like to thank Sandra Pariseau for administrative support. Additionally, we would like to thank Chris Bolick, Eileen Williams, and the staff at Marsh Botanical Gardens for their support in maintaining plant growth spaces. We would like to thank Dr. Shirin Bahmanyar for help with confocal microscopy. We would also like to thank Dr. Shirin Bahmanyar, Dr. Qingqing Wang, Christopher Adamchek, Harper Lowrey, and Lilijana Oliver for critical reading of the manuscript. We would like to thank Dr. Joel Greenwood for invaluable technical support for our automated luciferase imaging system. W.L., C.L., A.F., D.T., B.T., X.W., M.V., G.S., D.C. and J.M.G. designed the experiments. W.L., C.L., A.F., D.T., A.S., M.V., G.S. and W.Y. performed the experiments and experimental analyses. W.L., C.L., A.F., and J.M.G. wrote the article. This work was supported by the National Science Foundation (EAGER #1548538) and the National Institutes of Health (R35 GM128670) to J.M.G; and by the National Institutes of Health (T32 GM007499), the Gruber Foundation, and the National Science Foundation (GRFP DGE-1122492) to A.F. W.L. was supported by the Forest BH and Elizabeth DW Brown Fund Fellowship. DT was supported by the National Institutes of Health (T32GM007223-44).

## Figure Legends

**Figure S1.**
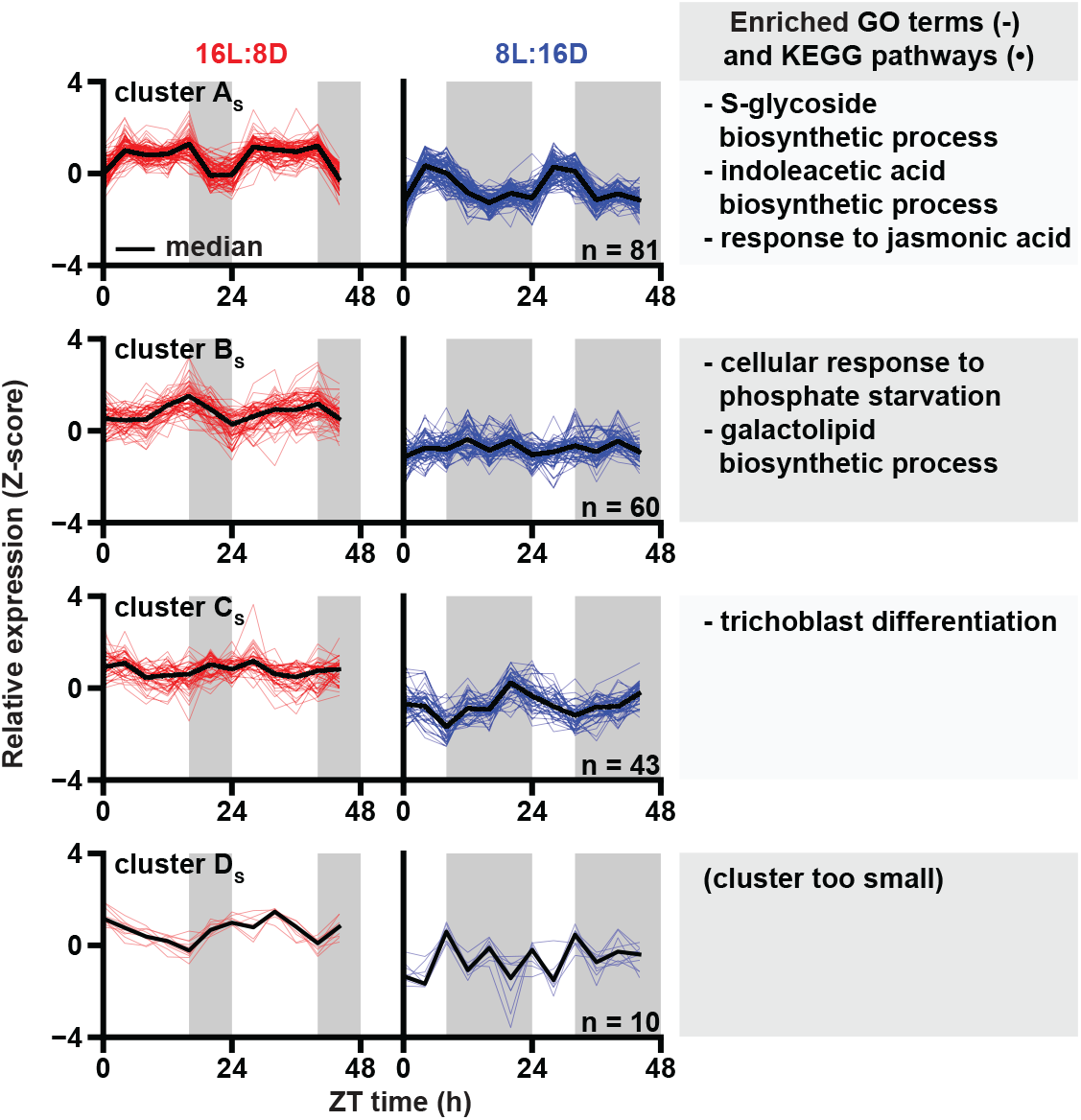
Summer gene expression pattern clusters. Normalized expression of 16L:8D-induced transcripts (rDEI_8L:16D/16L:8D_ < 0.5) grouped by k-means clustering (see Methods). 16L:8D (red) and 8L:16D (blue) expression patterns were transformed to Z score together for clustering to retain relative magnitude. Black lines indicate median expression level. Grey rectangles indicate the dark period of each photoperiod. The number of clusters is determined by the elbow method. Top enriched Gene Ontology (GO) and Kyoto Encyclopedia of Genes and Genomes (KEGG) pathways.

**Figure S2.**
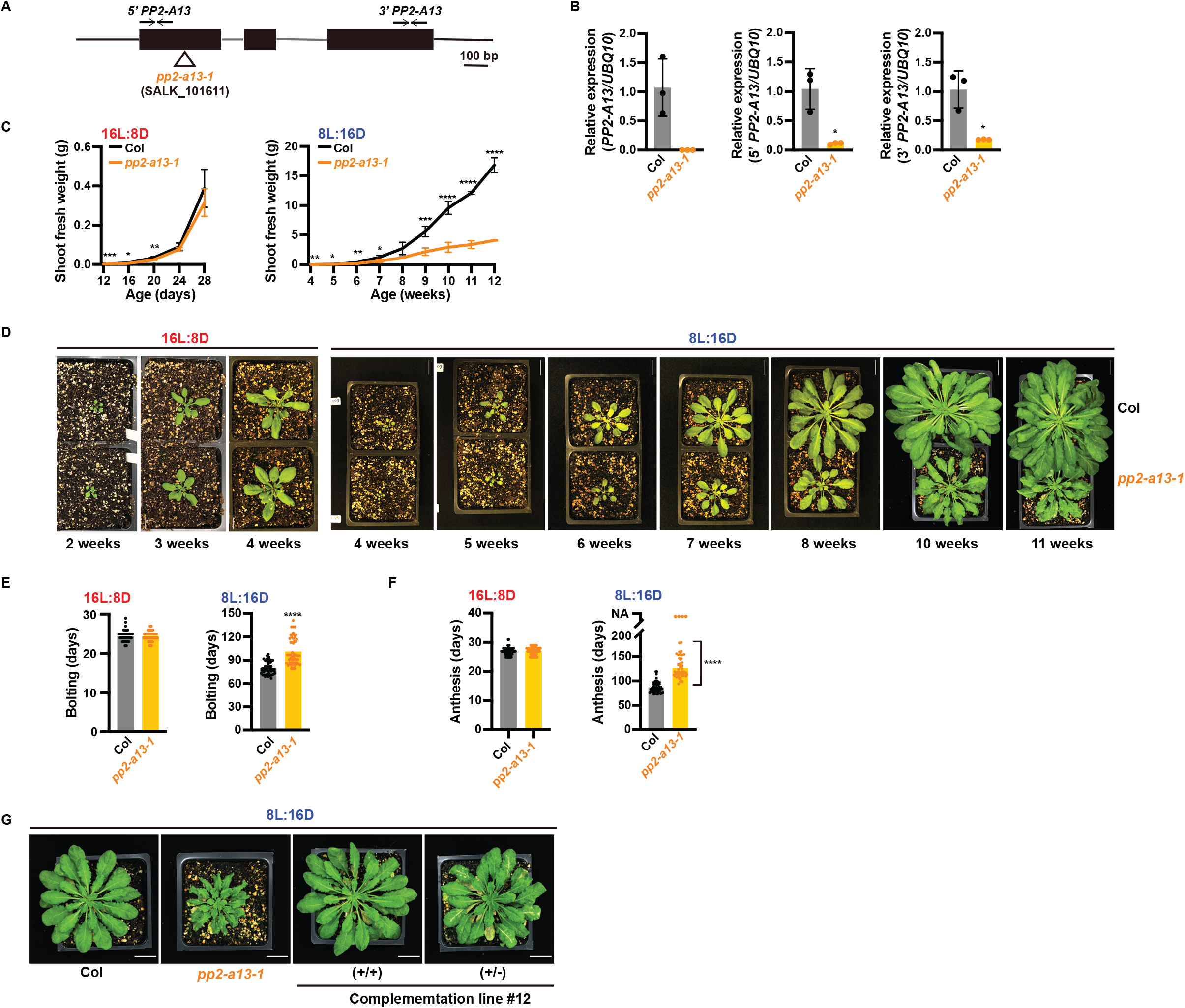
*PP2-A13* is needed for proper development and fitness in winter photoperiods. (A) Schematic shows the T-DNA insertion site in *PP2-A13*. Black boxes = exons; black lines = non-coding sequences. (B) qRT-PCR of full length, 5’ end, and 3’ end of the *PP2-A13* gene. Tissue was collected at ZT11 from 12-day-old plants grown in 8L:16D. *UBQ10* was used as internal control. n = 3 samples containing multiple seedlings. Error bar indicates SD. *, *p*<0.05 (Welch’s t-test). (C) Aerial fresh weight of wild-type (Col) and *pp2-a13-1* mutant plants grown in 16L:8D and 8L:16D. Error bar indicates SD. *, *p*≤ 0.05; **, *p*≤0.01; ***, *p*≤0.001; ****, *p*≤0.0001 (Welch’s t-test). (D) Representative images of wild-type (Col) and *pp2-a13-1* mutant plants at different time points prior to flowering. Plants grown in 16L:8D and 8L:16D. Scale bar = 2 cm in 16L:8D and 3 cm in 8L:16D. (E) Number of days until appearance of 1 cm long bolt for wild-type (Col) and *pp2-a13-1* mutant plants grown in 16L:8D and 8L:16D. *n*= 52-60. ****, *p*<0.0001 (Welch’s t-test). (F) Number of days until anthesis of the first flower for wild-type (Col) and *pp2-a13-1* mutant plants grown in 16L:8D and 8L:16D. n = 52-60. Welch’s t-test was performed on values excluding the four non-anthesed plants. ****, *p*<0.0001. (G) Segregating progeny from *PP2-A13*_*promoter*_::*gPP2-A13* complementation lines in the *pp2-a13-1* mutant background. +/+ and +/− indicate homozygous and hemizygous for the transgene, respectively. Images were taken of 9-week-old plants grown in 8L:16D. Scale bar = 3 cm.

**Figure S3.**
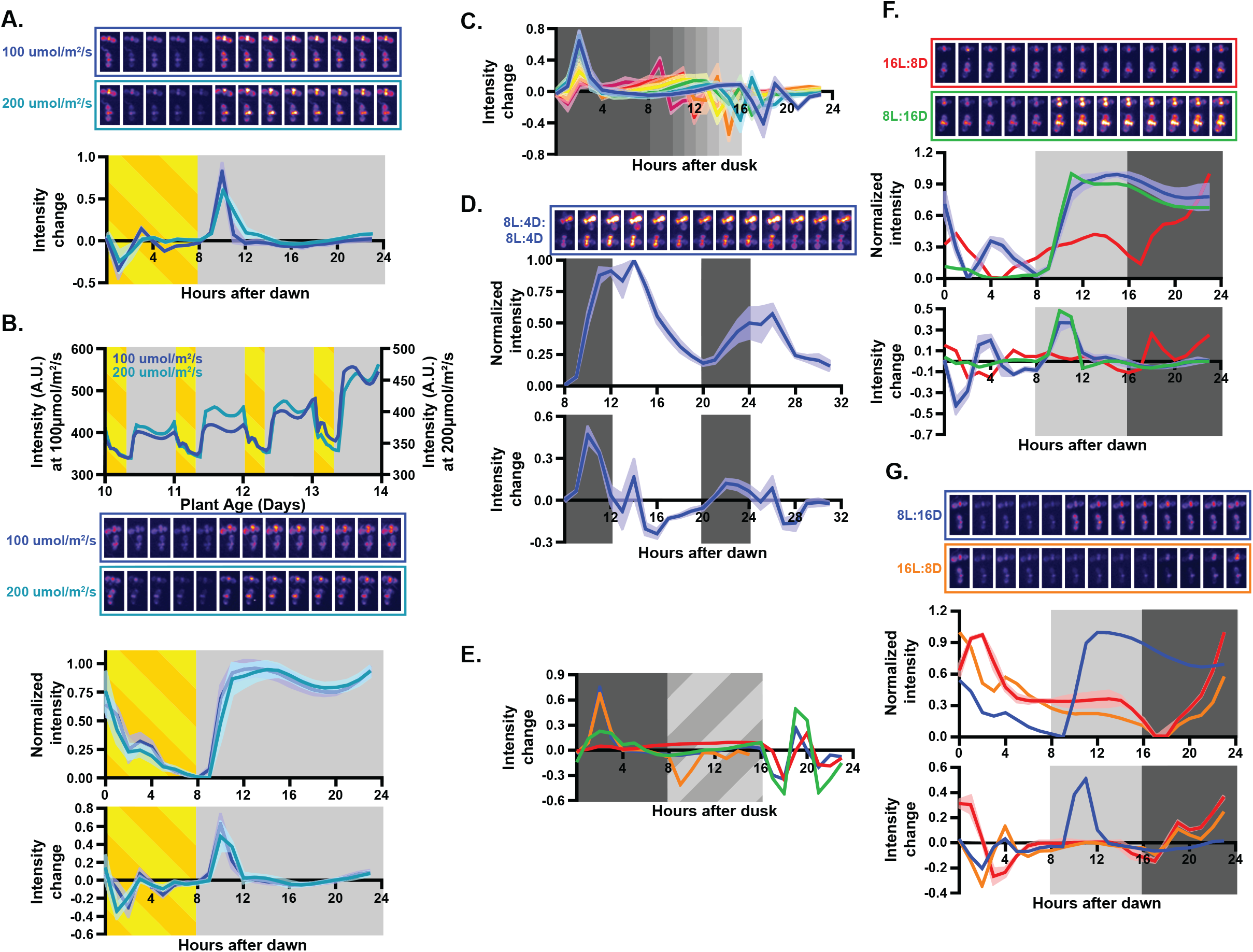
*PP2-A13* expression is controlled by photoperiod. (A) Representative images and intensity changes for data presented in figure 4E. (B) *PP2-A13*_*promoter*_::*Luciferase* expression in plants grown under short day conditions with either 100 μM m^−2^ s^−1^ (blue) or 200 μM m^−2^ s^−1^ (teal) white light. (C) Intensity change calculations for data presented in figure 4F. (D) Representative images, normalized traces, and intensity changes for traces presented in figure 4H. (E) Intensity changes for data presented in figure 4I. (F) Representative images, normalized intensity, and intensity changes for figure 4J. (G) Representative images, normalized intensity, and intensity changes for figure 4K.

**Figure S4.**
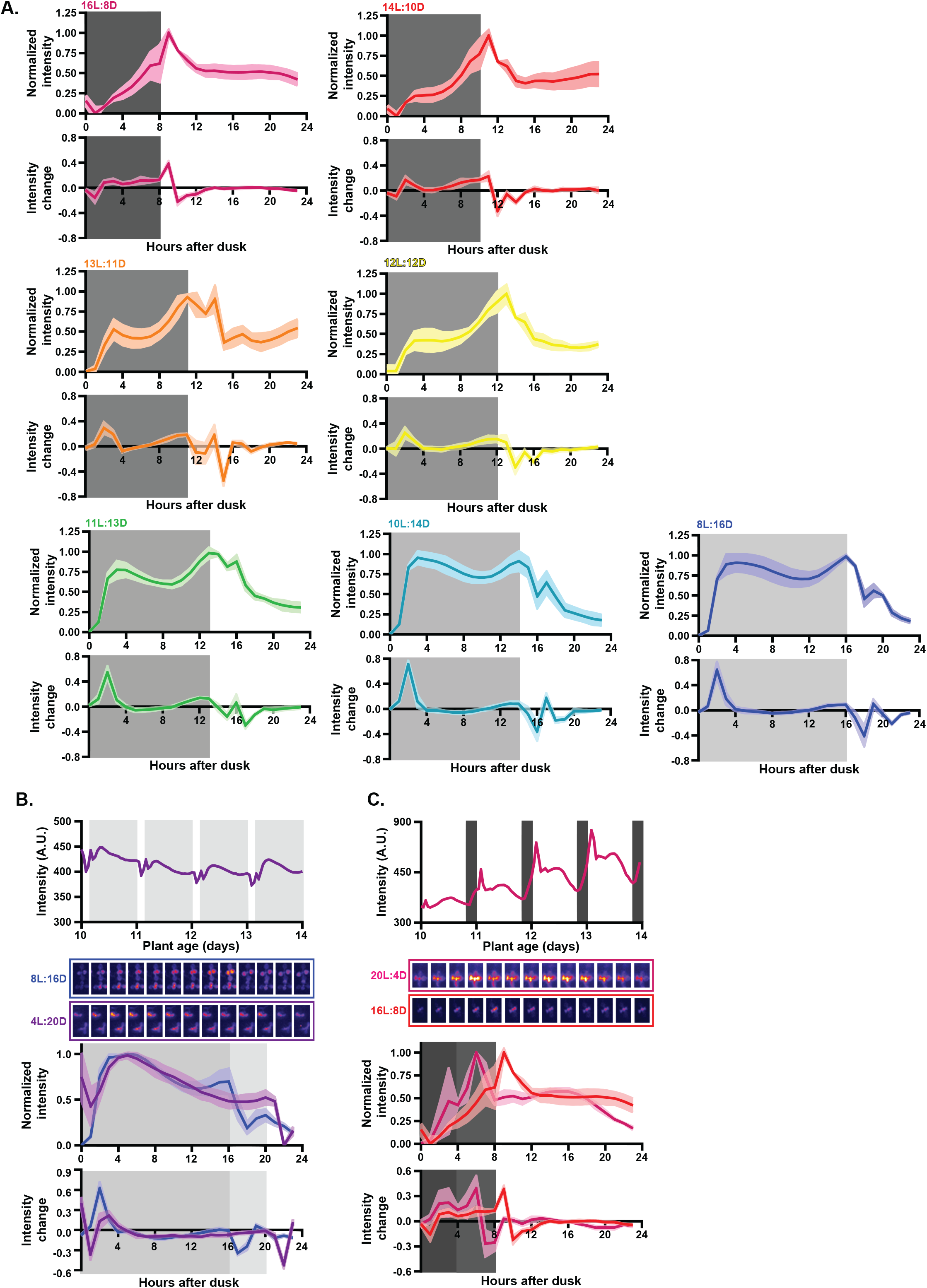
*PP2-A13* critical photoperiod. (A) Data is same as in Figure 4F except plotted independently for clarity. (**B**) *PP2-A13*_*promoter*_::*Luciferase* expression in plants grown under 4L:20D conditions (purple) and 20L:4D conditions (magenta).

**Figure S5.**
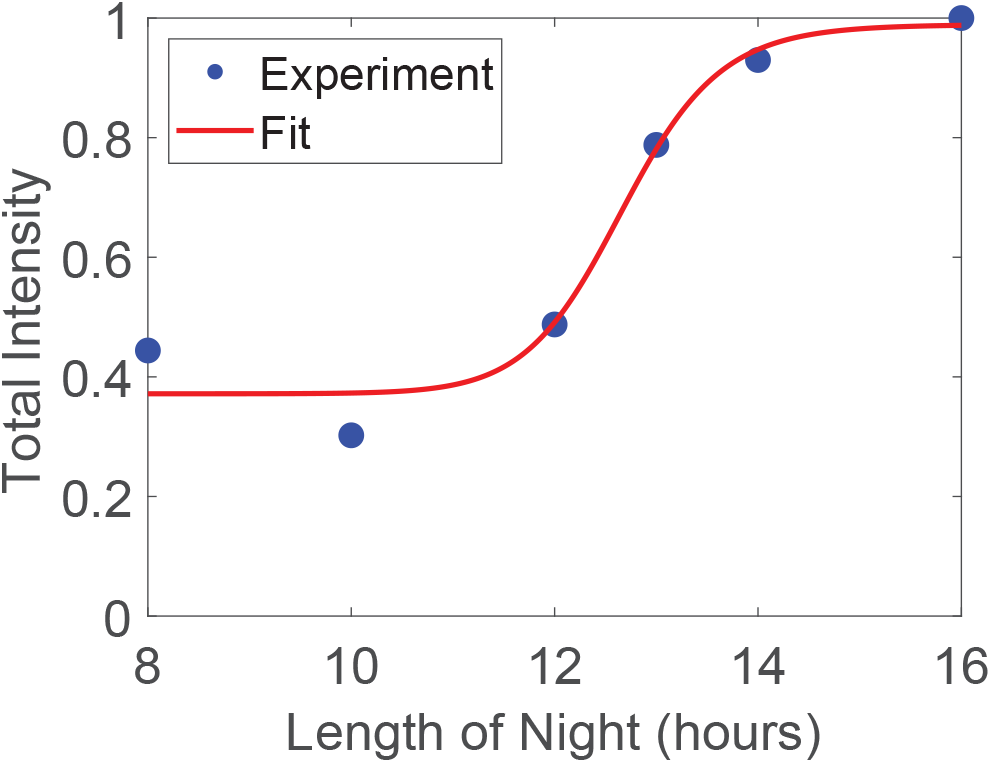
Curve fit for estimated yearly expression of *PP2-A13*_*promoter*_::*Luciferase*. Approximately sigmoidal fit to the total, normalized intensity of *PP2-A13*_*promoter*_::*luciferase* in a day. Blue points are the experimental points from the 6 conditions in figure S4. Red line is the approximately sigmoidal fit.

**Figure S6.**
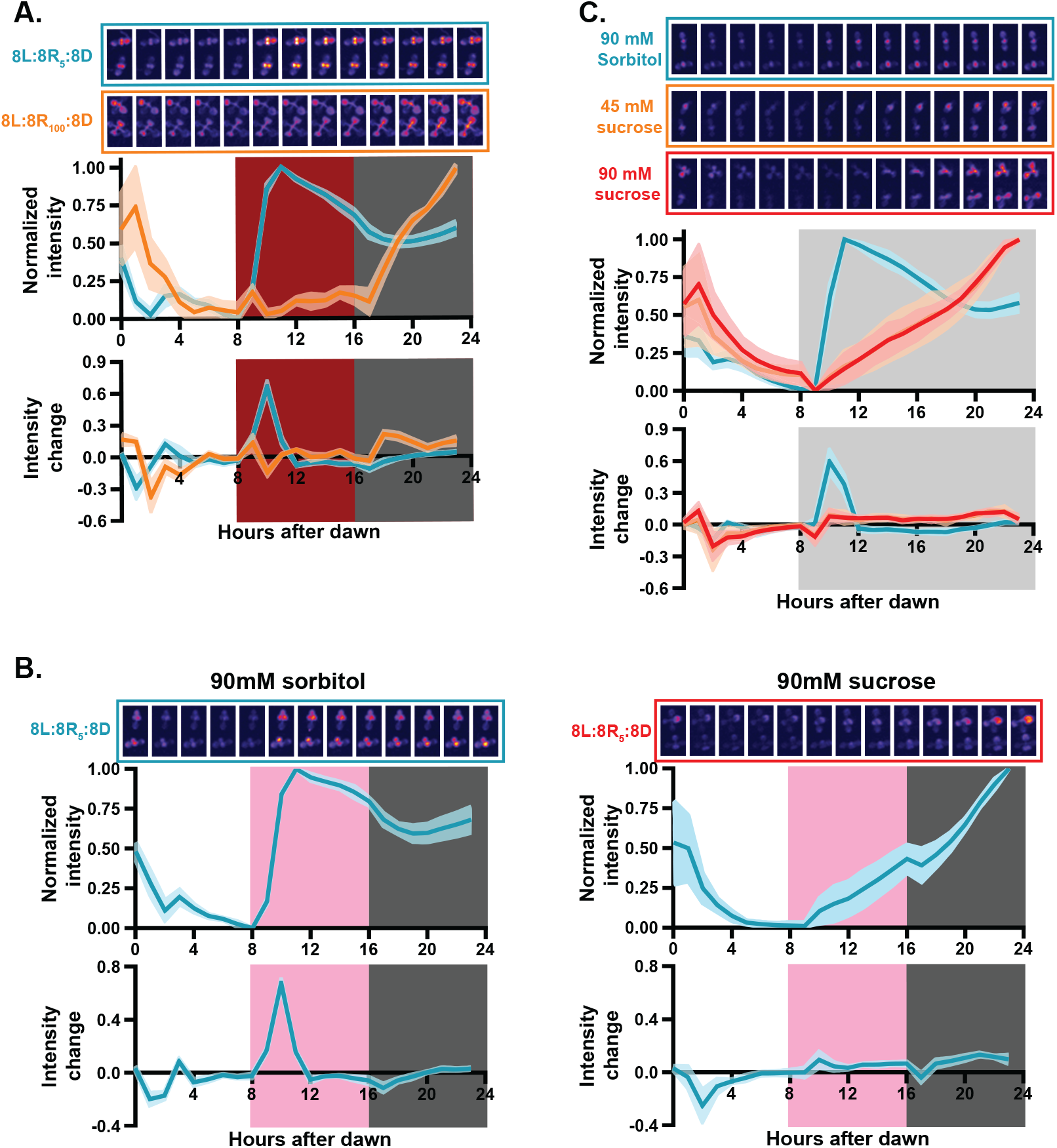
The photosynthetic apparatus senses darkness for winter photoperiod time measurement. (A) Representative images, normalized traces, and intensity changes for traces presented in figure 6A. (B) Representative images, normalized traces, and intensity changes for traces presented in figure 6B. (C) Representative images, normalized traces, and intensity changes for traces presented in figure 6C.

**Figure S7.**
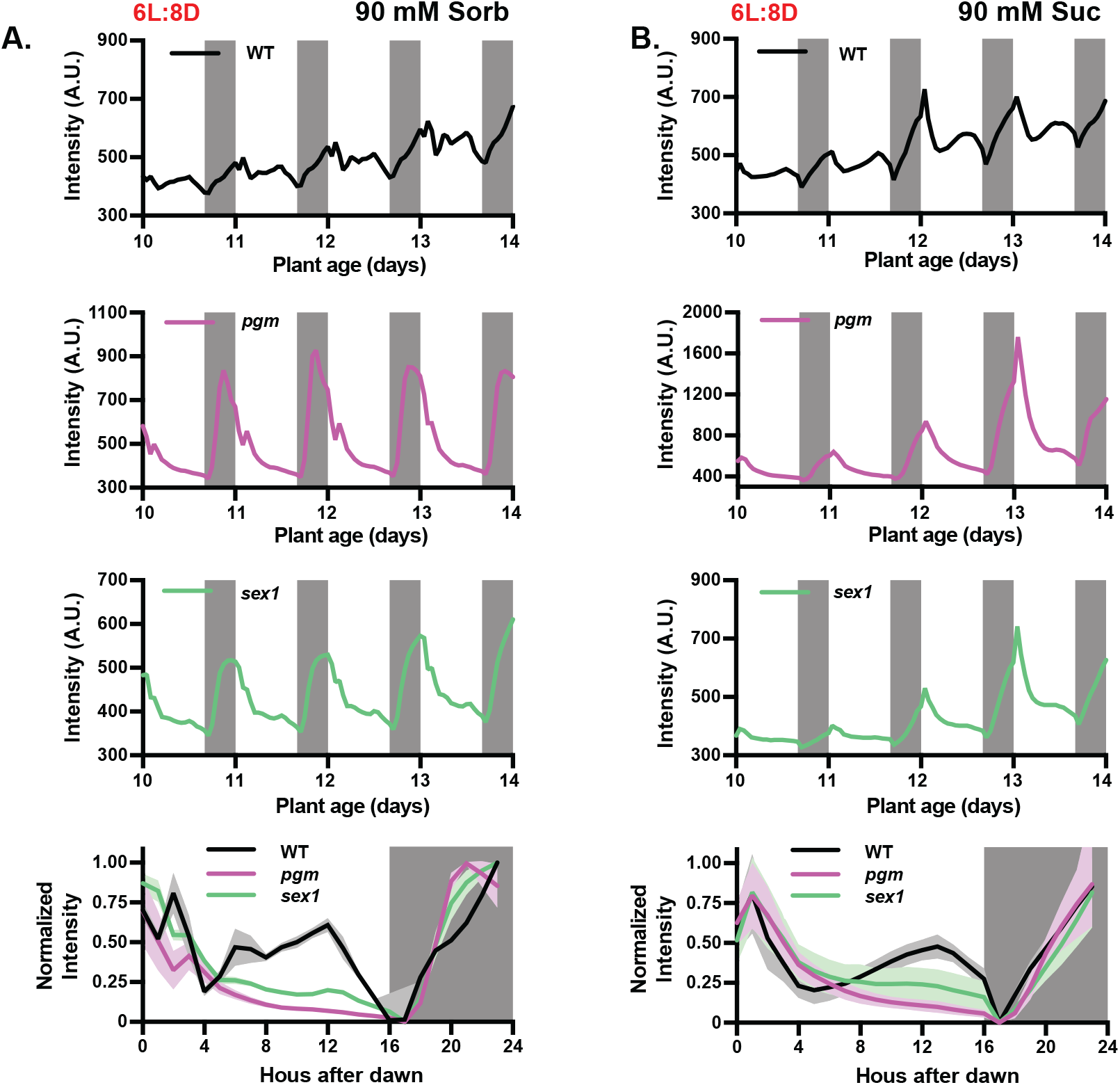
A metabolic coincidence mechanism controls winter photoperiod gene expression. (A-B) Traces and normalized trace data from plants grown in 16L:8D with (A) 90mM sorbitol or (B) 90mM sucrose.

Table S1: Relative daily expression integral (rDEI) of the probes of ATH1 microarray.

Table S2: Description and enriched annotations of 8L:16D-induced clusters.

Table S3: Description and enriched annotations of 16L:8D-induced clusters.

Table S4: Description and enriched annotations of clusters identified by hierarchical clustering. Table S5: Primers used in this study.

## Materials and Methods

### Plant materials and growth conditions

The *PP2-A13*_*promoter*_::*luciferase* transgenic line was generated in this study as described in section “plasmid construction”. The *PP2-A13* complementation line was generated by transformation of agrobacteria GV3101 harboring *PP2-A13*_*promoter*_::*gPP2-A13* construct into the *pp2-a13-1* background. The transgenic lines were selected by hygromycin and genotyping. The Arabidopsis seeds of Col-0, *pp2-a13-1* (SALK_101611), *pgm-1* (CS210), *sex1-1* (CS3093), *co-9* (CS870084), *atg5-1* (CS39993), and *atg7-2* (CS69859) were obtained from ABRC. *pp2-a13-1* was also crossed to *atg5-1*, *atg7-2*, and *pgm* mutants and the double mutants were identified by genotyping. *PP2-A13*_*promoter*_::*luciferase* transgenic line was crossed to *co-9*, *pgm*, and *sex1* mutant and the homozygous lines were identified by genotyping and bioluminescence imaging. The *pgm-1* allele was genotyped as described by (Veley, et al., 2012). The *sex1-1* allele was genotyped by PCR followed by *StyI* digestion (WT = 387 bp + 607 bp). The primers used for genotyping are listed in table S5.

Regarding samples for qRT-PCR assays, seeds from Arabidopsis Col-0 or the indicated mutant were sown on filter paper soaked with 0.5X Murashige and Skoog agar plates (pH 5.7) and stratified at 4°C for 2 days in the dark. Afterwards, the plates were transferred to a growth chamber at 22°C illuminated by white fluorescent lamps at 150 μmol m^2^ s^−1^ under photoperiod of 16L:8D, 12L:12D, or 8L:16D for the indicated duration. Specifically, for figure 2A, seeds were given 24 hours for germination and the seedlings were harvested on the thirteenth day after germination. Triplicates were collected every 4 hours starting at ZT0. For the ZT0 time point, collection took place 5 minutes before dawn. For the dusk time point of the respective photoperiod, collection took place in the light. Whole seedlings were snap-frozen with liquid nitrogen. For soil-grown plants, after two days stratification, seeds were germinated and grown in Fafard-2 mix at 22 °C under 16L:8D or 8L:16D.

### Plasmid construction

For the *PP2-A13* complementation plasmids, the *PP2-A13*_*promoter*_::*gPP2-A13* fragment was generated from PCR using Col-0 genomic DNA as template and inserted into pENTR/D-TOPO vectors (Invitrogen, cat. # K240020) then transferred into pGWB16 destination vectors using LR recombination(Nakagawa, et al., 2007).

To generate the *PP2-A13*_*promoter*_::*LUC* construct, a 2233 bp promoter sequence upstream the *PP2-A13* coding sequence was obtained by PCR and inserted into pENTR vector and then transferred into the pFLASH destination vectors to drive the luciferase(Gendron, et al., 2012).

To generate the *PP2-A13*_*promoter*_::*GUS* construct, the 2233 bp promoter sequence was subcloned from entry vector pENTR*-PP2-A13pro* to destination vector pMDC164 by LR recombination (Curtis and Grossniklaus, 2003). The primers used for cloning are listed in table S5.

### Luciferase Imaging and Analysis

*PP2-A13*_*promoter*_::*Luciferase* and *DIN6*_*promoter*_::*Luciferase* seeds were surface sterilized for 20 minutes in 70% ethanol and 0.01% Triton X-100 then sown on freshly poured ½ MS plates (2.15 g/L Murashige and Skoog medium, pH 5.7, Cassion Laboratories, cat. # MSP01 and 0.8% bacteriological agar, AmericanBio cat. # AB01185) without sucrose. Seeds were stratified in the dark for two days at 4°C then transferred into 22°C, 12L/12D conditions for seven days. Twenty seven-day old seedlings were transferred onto freshly poured 100 mm square ½ MS plates with and without added sugars as indicated for a given experiment, in a 10×10 grid. Seedlings were then treated with5 mM D-luciferin (Cayman Chemical Company, cat. # 115144-35-9) dissolved in 0.01% TritonX-100, and imaged at 22°C under the indicated conditions. Under light conditions, lights were on for 52 minutes of every hour: the lights are off for two minutes prior to a five minute exposure collected on an Andor iKon-M CCD camera, and then remain off for one minute following the exposure. During the dark period, images were taken during the same five minute time period. Light was provided by two LED light panels (Heliospectra L1) with light fluence rate of 100-150 μmol m^−2^ s^−1^, unless otherwise indicated. The CCD camera was controlled using Micromanager, using the following settings: binning of 2, pre-amp gain of 2, and a 0.05 MHz readout mode(Edelstein, et al., 2014). Using this setup, up to 400 seedlings are simultaneously imaged across four plates. Images are acquired each hour for approximately six and a half days. Data was collected for the entire imaging period (the end of day 7 through the dawn of day 14) but only the data collected between days 10 and 14 of plant growth are presented in figures and used for analyses. The mean intensity of each seedling at each time point was calculated using ImageJ(Schneider, et al., 2012). These raw values are presented as raw trace plots.

### Normalization of luciferase imaging data

For normalization, the maximum and minimum expression values in a 25 hour period (defined as either one hour before dawn to the subsequent dawn or one hour before dusk to the subsequent dusk, as indicated for each experiment) were calculated. The minimum expression value was subtracted from each expression value, then this value was divided by the difference in expression between the maximum and minimum expression within that 24 hour period.

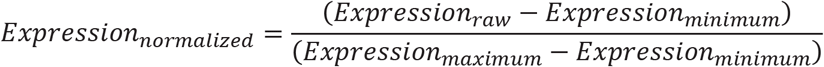

The mean and standard deviation of these normalized expression values were calculated for all days within an experiment of the same light conditions, unless otherwise indicated. Only the normalized expression values from dawn to dawn or dusk to dusk are plotted. The rate of change in expression was also calculated from the normalized expression values by calculating the difference between the expression at time *t* and the expression at time *t–1*. Because of the nature of this calculation, only 24 rate values are calculated. The mean and standard deviation of these rate values were calculated for all days within an experiment of the same light conditions, unless otherwise indicated.

### Estimation of yearly expression of *PP2-A13*_*promoter*_::*luciferase*

The total PP2-A13_promoter_::Luciferase intensity is first determined by taking the area under the curve, using the trapezoidal rule for numerical integration, for the six different light/dark conditions in figure S4. Since the plots in figure S4 are averaged over multiple days, a correction in the total *PP2-A13*_*promoter*_::*luciferase* intensity for the growth of the plant should be included. This is done by taking the intensity value at dusk and at 23 hours after dusk, connecting these points with a straight line, evaluating the resulting area under the curve (area of a triangle), then subtracting the total area under the curve by that triangular area. The area correction helps diminish the effects of plant growth. These corrected areas are then divided by the largest value (the 8L:16D condition) to obtain the normalized *PP2-A13*_*promoter*_::*luciferase* intensity. The normalized intensities are then fit with an approximately sigmoid function

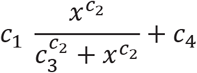

The built in non-linear data fitting tool in Xmgrace was used to determine the best fit parameters to the data are *c*_1_ = 0.62, *c*_2_ = 26.27, *c*_3_ = 12.67, and *c*_4_ = 0.37.

Using the sigmoidal fit from figure S5, the expression of *PP2-A13*_*promoter*_::*luciferase* over the course of a year is estimated. Since *Arabidopsis* Columbia ecotype was first isolated in Landsberg, Germany (https://peerj.com/preprints/26931v5/) (latitude, ~4g° N), the length of the night for each day in 2019 in Landsberg, Germany (https://www.timeanddate.com/sun/germany/landsberg-am-lech) was used to estimate the daily normalized expression of *PP2-A13*_*promoter*_::*luciferase*.

### qRT-PCR

For qRT-PCR experiments, RNA extraction was performed with two different methods. For figures 3A and 5A, total RNA was extracted from Arabidopsis seedlings grown in indicated conditions using TRIzol™ reagent (ThermoFisher, cat. #15596026); for the remaining figures 4B, 4D, 4E, 5D, 5E, and S2D, extraction was performed with RNeasy Plant Mini Kit (QIAGEN cat. # 74904). In both methods, the resulted RNA was subsequently treated with DNase (QIAGEN, cat. # 79254). The subsequent reverse-transcription and conditions for qRT-PCR reactions were described previously with minor modifications(Lee, et al., 2018). Four hundred nanograms of total RNA were used for reverse-transcription using iScript™ Reverse Transcription Supermix for RT-qPCR (Bio-Rad, cat. # 1708841). iTaq Universal SYBR Green Supermix was used for qRT-PCR reaction (Bio-Rad, cat. # 1725121). *IPP2* (AT3G02780) or *UBQ10* (AT4G05320) was used as an internal control as indicated. The relative expression represents means of 2^(-KCT)^ from three biological replicates, in which KCT = (CT of Gene of Interest – CT of internal control). The primers are listed in Table S5.

### Clustering analysis

The time-course microarray dataset was downloaded from the DIURNAL database (ftp://www.mocklerlab.org/diurnal)(Michael, et al., 2008; Mockler, et al., 2007). Relative daily expression integral for a transcript was calculated as: (sum of expression values in the DIURNAL “shortday” 8L:16D condition) / (sum of expression values in the DIURNAL “longday” 16L:8D condition). For the k-means clustering by both 16L:8D and 8L:16D expression values (Fig.1B), we performed log2-transformation followed by Z-score transformation in a gene-wise manner across both 16L:8D and 8L:16D expression values. We performed k-means clustering with the ‘kmeans’ function from scikit-learn python package(Pedregosa, et al., 2011) and determined the number of clusters using the elbow method with inertia.

For the hierarchical clustering analysis (Fig.1C), we performed log2-transformation of the data followed by Z-score transformation in a gene-wise manner separately for each time course to obtain the pattern. Principal components amounting to just above 90% of the total variance were used for clustering using the ‘factoextra’ R package (Alboukadel Kassambara and Fabian Mundt (2020). factoextra: Extract and Visualize the Results of Multivariate Data Analyses. R package version 1.0.7. https://CRAN.R-project.org/package=factoextra). Gene-wise Pearson correlation was used as similarity measure for hierarchical clustering using the R ‘hclust’ function with average linkage. The ‘cutreeDynamic’ function from the ‘dynamicTreeCut’ R package(Langfelder, et al., 2008) was used to identify clusters from the dendrogram, with the parameters: method=“hybrid”, minClusterSize=50, deepSplit=1, pamStage=FALSE. For figure 1D, clusters of strongly photoperiodic expression were identified by testing the mean log2(rDEI_8L:16D/16L:8D_) of the cluster against zero using the one-sample Wilcoxon signed rank test. All three identified clusters with −log10(adjusted *p*-value) > 20 (Bonferroni correction) were 8L:16D-induced. All code used for clustering analysis are provided in the supplementary materials.

### Functional enrichment analysis

Only clusters that have at least 40 transcripts were tested for enrichment of functional annotations. Enrichment analysis of Gene Ontology (GO) terms and Kyoto Encyclopedia of Genes and Genomes (KEGG) pathways was performed with the R package ‘clusterProfiler’, using the enrichGO function and the enrichKEGG function with the parameters: pAdjustMethod = “BH”, pvalueCutoff = 0.05, qvalueCutoff = 0.05, respectively(Yu, et al., 2012; Hvidsten, et al., 2001; Kanehisa and Goto, 2000; Ogata, et al., 1998). Highly similar GO terms were merged with the ‘simplify’ function with the parameters: cutoff = 0.5, measure = ‘Wong’, by=’p.adjust’. Since redundant annotations were still present after merging, notable annotations were manually selected for figure 1B. The full list of annotations is available in the Supplementary materials.

### GUS histochemical analysis

For GUS assay, the *PP2-A13*_*promoter*_::*GUS* transgenic plant was grown in 12L:12D for 12 days and then transferred to 8L:16D for 3 more days. The plant was freshly harvested and stained at 37 °C over night with 2 mM 5-bromo-4-chloro-3-indolyl-beta-D-glucuronic acid (X-glu) in 100 mM potassium phosphate buffer, pH 7.0, containing 0.1% (v/v) Triton X-100, 1 mM K3Fe(CN)6 and 10 mM EDTA. Tissues were cleared before observation by washing with 75% (v/v) ethanol.

### Subcellular localization

For subcellular localization studies, the coding sequences of the *PP2-A13* gene were recombined into pGW-GFP vector which harbors an in-frame C-terminal GFP and is driven by the *Cauliflower mosaic virus* (*CaMV*) *35S* promoter. The *35S::PP2-A13-GFP* construct was co-transformed with *35S::mCherry-VirD2NLS* as a nuclear marker (Citovsky, et al., 2006). Arabidopsis protoplast transfection was performed as previously described (Yoo, et al., 2007) and the subcellular localization of the fluorescent-tagged protein was detected with a Nikon ECLIPSE Ti confocal microscope system.

### Immunoblotting

For immunoblot analysis, WT and *pp2-a13-1* mutant plants were ground in liquid nitrogen. Crude proteins were extracted with SII buffer (100 mM sodium phosphate, pH 8, 150 mM NaCl, 5 mM EDTA, and 0.1% [v/v] Triton X-100) with cOmplete EDTA-free Protease Inhibitor Cocktail (Roche, catalog no. 11873580001) and 1 mM phenylmethylsulfonyl fluoride. Protein concentration was quantified with a Pierce BCA Protein Assay Kit (Thermo Fisher Scientific, catalog no. 23225). Approximately 50 μg of total protein was loaded and separated on 12% (w/v) SDS-PAGE for immunoblot analyses. ATG8a and actin protein levels were detected with anti-ATG8a antibody (1:1000; abcam, ab77003) and anti-actin antibody (1:3000; Millipore-Sigma, SAB4301137).

### Glycoprotein staining

For glycoprotein staining, the procedure of protein extraction, quantification, and separation are the same as the procedure in section “Immunoblotting”. The glycoproteins in polyacrylamide gel was detected with Pierce Glycoprotein Staining Kit (catalog no. 24562) according to the manufacture’s procedure.

## References

An, H., Roussot, C., Suarez-Lopez, P., Corbesier, L., Vincent, C., Pineiro, M., Hepworth, S., Mouradov, A., Justin, S., Turnbull, C., et al. (2004). CONSTANS acts in the phloem to regulate a systemic signal that induces photoperiodic flowering of Arabidopsis. Development (Cambridge, England) 131, 3615–26.

Azeez, A., and Sane, A.P. (2015). Photoperiodic growth control in perennial trees. Plant signaling & behavior 10, e1087631.

Bohlenius, H., Huang, T., Charbonnel-Campaa, L., Brunner, A.M., Jansson, S., Strauss, S.H., and Nilsson, O. (2006). CO/FT regulatory module controls timing of flowering and seasonal growth cessation in trees. Science (New York, N.Y.) 312, 1040–3.

Bouche, F., Woods, D.P., and Amasino, R.M. (2017). Winter Memory throughout the Plant Kingdom: Different Paths to Flowering. Plant physiology 173, 27–35.

Bradshaw, W.E., and Holzapfel, C.M. (2010). What season is it anyway? Circadian tracking vs. photoperiodic anticipation in insects. J Biol Rhythms 25, 155–65.

Bunning, E. (1969). Common features of photoperiodism in plants and animals. Photochem Photobiol 9, 219–28.

Caspar, T., Lin, T.P., Kakefuda, G., Benbow, L., Preiss, J., and Somerville, C. (1991). Mutants of Arabidopsis with altered regulation of starch degradation. Plant physiology 95, 1181–8.

Castillejo, C., and Pelaz, S. (2008). The balance between CONSTANS and TEMPRANILLO activities determines FT expression to trigger flowering. Current biology : CB 18, 1338–43.

Cornah, J.E., Germain, V., Ward, J.L., Beale, M.H., and Smith, S.M. (2004). Lipid utilization, gluconeogenesis, and seedling growth in Arabidopsis mutants lacking the glyoxylate cycle enzyme malate synthase. The Journal of biological chemistry 279, 42916–23.

Cubas, P. (2020). Plant Seasonal Growth: How Perennial Plants Sense That Winter Is Coming. Current biology : CB 30, R21–R23.

Diez, J.M., Ibanez, I., Silander, J.A., Jr., Primack, R., Higuchi, H., Kobori, H., Sen, A., and James, T.Y. (2014). Beyond seasonal climate: statistical estimation of phenological responses to weather. Ecol Appl 24, 1793–802.

Dinant, S., Clark, A.M., Zhu, Y., Vilaine, F., Palauqui, J.C., Kusiak, C., and Thompson, G.A. (2003). Diversity of the superfamily of phloem lectins (phloem protein 2) in angiosperms. Plant physiology 131, 114–28.

Eimert, K., Wang, S.M., Lue, W.I., and Chen, J. (1995). Monogenic Recessive Mutations Causing Both Late Floral Initiation and Excess Starch Accumulation in Arabidopsis. The Plant cell 7, 1703–1712.

Feke, A., Liu, W., Hong, J., Li, M.W., Lee, C.M., Zhou, E.K., and Gendron, J.M. (2019). Decoys provide a scalable platform for the identification of plant E3 ubiquitin ligases that regulate circadian function. Elife 8.

Feke, A.M., Hong, J., Liu, W., and Gendron, J.M. (2020). A Decoy Library Uncovers U-Box E3 Ubiquitin Ligases That Regulate Flowering Time in Arabidopsis. Genetics 215, 699–712.

Fournier-Level, A., Perry, E.O., Wang, J.A., Braun, P.T., Migneault, A., Cooper, M.D., Metcalf, C.J., and Schmitt, J. (2016). Predicting the evolutionary dynamics of seasonal adaptation to novel climates in Arabidopsis thaliana. Proceedings of the National Academy of Sciences of the United States of America 113, E2812–21.

Foyer, C.H. (2018). Reactive oxygen species, oxidative signaling and the regulation of photosynthesis. Environ Exp Bot 154, 134–142.

Garbazza, C., and Benedetti, F. (2018). Genetic Factors Affecting Seasonality, Mood, and the Circadian Clock. Front Endocrinol (Lausanne) 9, 481.

Han, C., Ren, C., Zhi, T., Zhou, Z., Liu, Y., Chen, F., Peng, W., and Xie, D. (2013). Disruption of fumarylacetoacetate hydrolase causes spontaneous cell death under short-day conditions in Arabidopsis. Plant physiology 162, 1956–64.

Henderson, I.R., Shindo, C., and Dean, C. (2003). The need for winter in the switch to flowering. Annu Rev Genet 37, 371–92.

Hvidsten, T.R., Komorowski, J., Sandvik, A.K., and Laegreid, A. (2001). Predicting gene function from gene expressions and ontologies. Pac Symp Biocomput, 299–310.

Inoue, S., Dang, Q.L., Man, R., and Tedla, B. (2020). Photoperiod and CO2 elevation influence morphological and physiological responses to drought in trembling aspen: implications for climate change-induced migration. Tree Physiol 40, 917–927.

Izumi, M., Hidema, J., Makino, A., and Ishida, H. (2013). Autophagy contributes to nighttime energy availability for growth in Arabidopsis. Plant physiology 161, 1682–93.

Jang, S., Marchal, V., Panigrahi, K.C., Wenkel, S., Soppe, W., Deng, X.W., Valverde, F., and Coupland, G. (2008). Arabidopsis COP1 shapes the temporal pattern of CO accumulation conferring a photoperiodic flowering response. EMBO J 27, 1277–88.

Johansson, M., and Staiger, D. (2014). SRR1 is essential to repress flowering in non-inductive conditions in Arabidopsis thaliana. Journal of experimental botany 65, 5811–22.

Kanehisa, M., and Goto, S. (2000). KEGG: kyoto encyclopedia of genes and genomes. Nucleic Acids Res 28, 27–30.

Kardailsky, I., Shukla, V.K., Ahn, J.H., Dagenais, N., Christensen, S.K., Nguyen, J.T., Chory, J., Harrison, M.J., and Weigel, D. (1999). Activation tagging of the floral inducer FT. Science (New York, N.Y.) 286, 1962–5.

Kim, J.A., Kim, H.S., Choi, S.H., Jang, J.Y., Jeong, M.J., and Lee, S.I. (2017). The Importance of the Circadian Clock in Regulating Plant Metabolism. Int J Mol Sci 18.

Kreyling, J. (2010). Winter climate change: a critical factor for temperate vegetation performance. Ecology 91, 1939–48.

Lee C-M, F., A, Adamchek C, Webb K, Pruneda-Paz J, Bennett EJ, Kay SA, Gendron JM (2017). Decoys reveal the genetic and biochemical roles of redundant plant E3 ubiquitin ligases. biorXiv.

Lee, C.M., Feke, A., Li, M.W., Adamchek, C., Webb, K., Pruneda-Paz, J., Bennett, E.J., Kay, S.A., and Gendron, J.M. (2018). Decoys Untangle Complicated Redundancy and Reveal Targets of Circadian Clock F-Box Proteins. Plant physiology 177, 1170–1186.

Lumsden, P.J., and Millar, A.J. (1998). Biological rhythms and photoperiodism in plants (Oxford Washington, D.C. Herndon, VA: Bios Scientific Publishers; Bios Scientific Publishers distributor)

Mengin, V., Pyl, E.T., Alexandre Moraes, T., Sulpice, R., Krohn, N., Encke, B., and Stitt, M. (2017). Photosynthate partitioning to starch in Arabidopsis thaliana is insensitive to light intensity but sensitive to photoperiod due to a restriction on growth in the light in short photoperiods. Plant, cell & environment 40, 2608–2627.

Michael, T.P., Mockler, T.C., Breton, G., McEntee, C., Byer, A., Trout, J.D., Hazen, S.P., Shen, R., Priest, H.D., Sullivan, C.M., et al. (2008). Network discovery pipeline elucidates conserved time-of-day-specific cis-regulatory modules. PLoS Genet 4, e14.

Millar, A.J., Carre, I.A., Strayer, C.A., Chua, N.H., and Kay, S.A. (1995a). Circadian clock mutants in Arabidopsis identified by luciferase imaging. Science (New York, N.Y.) 267, 1161–3.

Millar, A.J., Short, S.R., Chua, N.H., and Kay, S.A. (1992). A novel circadian phenotype based on firefly luciferase expression in transgenic plants. The Plant cell 4, 1075–87.

Millar, A.J., Straume, M., Chory, J., Chua, N.H., and Kay, S.A. (1995b). The regulation of circadian period by phototransduction pathways in Arabidopsis. Science (New York, N.Y.) 267, 1163–6.

Mockler, T.C., Michael, T.P., Priest, H.D., Shen, R., Sullivan, C.M., Givan, S.A., McEntee, C., Kay, S.A., and Chory, J. (2007). The DIURNAL project: DIURNAL and circadian expression profiling, model-based pattern matching, and promoter analysis. Cold Spring Harb Symp Quant Biol 72, 353–63.

Moraes, T.A., Mengin, V., Annunziata, M.G., Encke, B., Krohn, N., Hohne, M., and Stitt, M. (2019). Response of the Circadian Clock and Diel Starch Turnover to One Day of Low Light or Low CO2. Plant physiology 179, 1457–1478.

Nakane, Y., and Yoshimura, T. (2019). Photoperiodic Regulation of Reproduction in Vertebrates. Annu Rev Anim Biosci 7, 173–194.

Nelson, R.J., Denlinger, D.L., and Somers, D.E. (2010). Photoperiodism : the biological calendar (Oxford; New York: Oxford University Press)

Ogata, H., Goto, S., Fujibuchi, W., and Kanehisa, M. (1998). Computation with the KEGG pathway database. Biosystems 47, 119–28.

Oquist, G., and Huner, N.P. (2003). Photosynthesis of overwintering evergreen plants. Annu Rev Plant Biol 54, 329–55.

Phillips, A.R., Suttangkakul, A., and Vierstra, R.D. (2008). The ATG12-conjugating enzyme ATG10 Is essential for autophagic vesicle formation in Arabidopsis thaliana. Genetics 178, 1339–53.

Putterill, J., Robson, F., Lee, K., Simon, R., and Coupland, G. (1995). The CONSTANS gene of Arabidopsis promotes flowering and encodes a protein showing similarities to zinc finger transcription factors. Cell 80, 847–57.

Roenneberg, T., and Merrow, M. (2001). Seasonality and photoperiodism in fungi. J Biol Rhythms 16, 403–14.

Saunders, D.S. (1997). Insect circadian rhythms and photoperiodism. Invert Neurosci 3, 155–64.

Saunders, D.S. (2005). Erwin Bunning and Tony Lees, two giants of chronobiology, and the problem of time measurement in insect photoperiodism. J Insect Physiol 51, 599–608.

Saunders, D.S. (2020). Dormancy, Diapause, and the Role of the Circadian System in Insect Photoperiodism. Annu Rev Entomol 65, 373–389.

Shim, J.S., and Imaizumi, T. (2015). Circadian clock and photoperiodic response in Arabidopsis: from seasonal flowering to redox homeostasis. Biochemistry 54, 157–70.

Song, Y.H., Shim, J.S., Kinmonth-Schultz, H.A., and Imaizumi, T. (2015). Photoperiodic flowering: time measurement mechanisms in leaves. Annu Rev Plant Biol 66, 441–64.

Stromme, C.B., Julkunen-Tiitto, R., Olsen, J.E., Nybakken, L., and Tognetti, R. (2017). High daytime temperature delays autumnal bud formation in Populus tremula under field conditions. Tree Physiol 37, 71–81.

Suarez-Lopez, P., Wheatley, K., Robson, F., Onouchi, H., Valverde, F., and Coupland, G. (2001). CONSTANS mediates between the circadian clock and the control of flowering in Arabidopsis. Nature 410, 1116–20.

Tan, Y., Merrow, M., and Roenneberg, T. (2004). Photoperiodism in Neurospora crassa. J Biol Rhythms 19, 135–43.

Valverde, F., Mouradov, A., Soppe, W., Ravenscroft, D., Samach, A., and Coupland, G. (2004). Photoreceptor regulation of CONSTANS protein in photoperiodic flowering. Science (New York, N.Y.) 303, 1003–6.

Vince-Prue, D. (1975). Photoperiodism in plants (London; New York: McGraw-Hill)

Vitasse, Y., Lenz, A., and Korner, C. (2014). The interaction between freezing tolerance and phenology in temperate deciduous trees. Front Plant Sci 5, 541.

Walker, W.H., 2nd, Melendez-Fernandez, O.H., Nelson, R.J., and Reiter, R.J. (2019). Global climate change and invariable photoperiods: A mismatch that jeopardizes animal fitness. Ecol Evol 9, 10044–10054.

Yanovsky, M.J., and Kay, S.A. (2002). Molecular basis of seasonal time measurement in Arabidopsis. Nature 419, 308–12.

Yoshida, Y., Adachi, E., Fukiya, K., Iwai, K., and Tanaka, K. (2005). Glycoprotein-specific ubiquitin ligases recognize N-glycans in unfolded substrates. EMBO Rep 6, 239–44.

Yoshida, Y., Mizushima, T., and Tanaka, K. (2019). Sugar-Recognizing Ubiquitin Ligases: Action Mechanisms and Physiology. Front Physiol 10, 104.

Yoshida, Y., Tokunaga, F., Chiba, T., Iwai, K., Tanaka, K., and Tai, T. (2003). Fbs2 is a new member of the E3 ubiquitin ligase family that recognizes sugar chains. The Journal of biological chemistry 278, 43877–84.

Zhi, T., Zhou, Z., Huang, Y., Han, C., Liu, Y., Zhu, Q., and Ren, C. (2016). Sugar suppresses cell death caused by disruption of fumarylacetoacetate hydrolase in Arabidopsis. Planta 244, 557–71.

## Materials and Methods References

Citovsky, V., Lee, L.Y., Vyas, S., Glick, E., Chen, M.H., Vainstein, A., Gafni, Y., Gelvin, S.B., and Tzfira, T. (2006). Subcellular localization of interacting proteins by bimolecular fluorescence complementation in planta. J Mol Biol 362, 1120–31.

Curtis, M.D., and Grossniklaus, U. (2003). A gateway cloning vector set for high-throughput functional analysis of genes in planta. Plant physiology 133, 462–9.

Edelstein, A.D., Tsuchida, M.A., Amodaj, N., Pinkard, H., Vale, R.D., and Stuurman, N. (2014). Advanced methods of microscope control using muManager software. J Biol Methods 1.

Gendron, J.M., Pruneda-Paz, J.L., Doherty, C.J., Gross, A.M., Kang, S.E., and Kay, S.A. (2012). Arabidopsis circadian clock protein, TOC1, is a DNA-binding transcription factor. Proceedings of the National Academy of Sciences of the United States of America 109, 3167–72.

Langfelder, P., Zhang, B., and Horvath, S. (2008). Defining clusters from a hierarchical cluster tree: the Dynamic Tree Cut package for R. Bioinformatics 24, 719–20.

Nakagawa, T., Kurose, T., Hino, T., Tanaka, K., Kawamukai, M., Niwa, Y., Toyooka, K., Matsuoka, K., Jinbo, T., and Kimura, T. (2007). Development of series of gateway binary vectors, pGWBs, for realizing efficient construction of fusion genes for plant transformation. J Biosci Bioeng 104, 34–41.

Pedregosa, F., Varoquaux, G., Gramfort, A., Michel, V., Thirion, B., Grisel, O., Blondel, M., Prettenhofer, P., Weiss, R., Dubourg, V., et al. (2011). Scikit-learn: Machine Learning in Python. J Mach Learn Res 12, 2825–2830.

Schneider, C.A., Rasband, W.S., and Eliceiri, K.W. (2012). NIH Image to ImageJ: 25 years of image analysis. Nat Methods 9, 671–5.

Veley, K.M., Marshburn, S., Clure, C.E., and Haswell, E.S. (2012). Mechanosensitive channels protect plastids from hypoosmotic stress during normal plant growth. Current biology : CB 22, 408–13.

Yoo, S.D., Cho, Y.H., and Sheen, J. (2007). Arabidopsis mesophyll protoplasts: a versatile cell system for transient gene expression analysis. Nat Protoc 2, 1565–72.

Yu, G., Wang, L.G., Han, Y., and He, Q.Y. (2012). clusterProfiler: an R package for comparing biological themes among gene clusters. OMICS 16, 284–7.

