## Supplementary Codes for "A metabolic coincidence mechanism controls winter photoperiodism in plants": 1_clustering.R

```
library(cluster)
library(FactoMineR)
library(factoextra)
library(dendextend)
library(dynamicTreeCut)
library(amap)


## Function to calculate Z-score and organise data
normalize <- function(raw_data, col_num_list){
  
  matrices <- list()
  for (k in 1:length(col_num_list)){
    columns <- col_num_list[[k]]
    series_name <- names(col_num_list)[k]
    mat_t <- as.matrix(read.csv(raw_data, stringsAsFactors=FALSE, row.names=1)[, columns])
    mat_tn <- t(scale(t(log(mat_t, 2)), center=TRUE, scale=TRUE))
    matrices[[series_name]] <- mat_tn
  }
  
  mat_combined <- do.call(cbind, matrices)
  
  res_pca <- prcomp(t(mat_tn), scale = TRUE)
  no_dim <- min(which((get_eig(res_pca)$cumulative.variance.percent>90)==TRUE))
  var_pca <- get_pca_var(res_pca)
  
  X <- var_pca$coord[, 1:no_dim]
  return(X)
}


## Function to perform hierarchical clustering and dynamic tree cutting
call_hclust <- function(condition, method, deepSplitNumber=1){
  
  X <- normalize(raw_data="diurnalValues.csv",
                 col_num_list=condition[[1]])
  
  res_hc <- hclust(Dist(X, method='correlation'), method=method)
  dend <- as.dendrogram(res_hc)
  res_cut <- cutreeDynamic(res_hc, distM = as.matrix(Dist(X, method='correlation')), method = "hybrid", minClusterSize=50, deepSplit=deepSplitNumber, pamStage=FALSE)
  names(res_cut) <- rownames(X)
  
  dump('res_hc', file=paste(c('hclust_', method, '_', condition[[2]], '.R'), collapse=''))
  dump('res_cut', file=paste(c('DTC_', method, '_', condition[[2]], '_d', as.character(deepSplitNumber), '_mc50_pamF.R'), collapse=''))
}


## List of list as input of call_hclust()
sdld <- list(columns=list('longday'=c(1:12),
                          'shortday'=c(13:24),
                          'ler_sd'=c(25:36)),
             name='sdld')


## Calling the function
call_hclust(condition=sdld, method='average', deepSplitNumber=1)


## Processing clustering data and performing GO term analysis
for (cond in c('sdld')){
  
  ## 01. Loading Data
  
  pipe_in <- './'
  pipe_out <- paste(c('output/', cond, '/'), collapse='')
  dir.create(pipe_out, showWarnings = FALSE)
  
  source('func_dendextend.R')
  dend <- as.dendrogram(dget(paste(c(pipe_in, 'hclust_average_', cond, '.R'), collapse='')))
  res_cut <- order_clusters_by_median(dend=dend,
                                      res_DTC=dget(paste(c(pipe_in, 'DTC_average_', cond, '_d1_mc50_pamF.R'), collapse='')))
  print(max(res_cut))
  
  ## 02. Extract the clusters and write them to files
  clusters <- convert_to_cluster_vector(res_cut)
  fileConn<-file(paste(c(pipe_out, 'clusters.txt'), collapse=''))
  writeLines(clusters, fileConn)
  close(fileConn)
  df_s <- read.csv('rDEI_ID.csv', row.names = 1)
  write.csv(make_cluster_descriptions(clusters=clusters, df_s=df_s),
            file=paste(c(pipe_out,'clusterDescription.csv'), collapse=''), row.names=FALSE)
  
  
  ## 03. GO term analysis
  source('func_ClusterProfiler.R')
  df_s2  <- read.csv('rDEI_ID.csv', stringsAsFactors=FALSE)
  call_enrichGO(clusters=clusters, df_s=df_s2, ont='BP', threshold=0, out_dir=paste(c(pipe_out, 'GO_BP/'), collapse=''))
  call_enrichGO(clusters=clusters, df_s=df_s2, ont='MF', threshold=0, out_dir=paste(c(pipe_out, 'GO_MF/'), collapse=''))
  call_enrichGO(clusters=clusters, df_s=df_s2, ont='CC', threshold=0, out_dir=paste(c(pipe_out, 'GO_CC/'), collapse=''))
  call_enrichKEGG(clusters=clusters, df_s=df_s2, threshold=0, out_dir=paste(c(pipe_out, 'KEGG/'), collapse=''))
}
```
