## Supplementary Codes for "A metabolic coincidence mechanism controls winter photoperiodism in plants": 2_makeDendrogram.R

```
library(cluster)
library(dendextend)
library(dynamicTreeCut)
library(colorspace)
library(circlize)
library(ComplexHeatmap)
library(scales)
library(svglite)


cond <-'sdld'
coloring_cond2 <- 'sdld'


## 01. Loading Data
pipe_in <- './'
pipe_out <- 'output/'
dir.create(pipe_out, showWarnings=FALSE)

source('func_dendextend.R')
dend <- as.dendrogram(dget(paste(c(pipe_in, 'hclust_average_', cond, '.R'), collapse='')))
res_cut <- order_clusters_by_median(dend=dend,
                                    res_DTC=dget(paste(c(pipe_in, 'DTC_average_', cond, '_d1_mc50_pamF.R'), collapse='')))


## 02. The plotting function
plot_dendrogram <- function(dend, df_s, coloring_cond,
                            color1='#0000FF', color2='#0F99B2', color3='#AAAAFF',
                            color1_clusters, color2_clusters, color3_clusters,
                            clusters_to_label,
                            filename){
  
  col_fun<-colorRamp2(c(-600, -50, 0, 50, 600), 
                      c("#FF8000", "#FF8000", "#000000", "#0F99B2", "#0F99B2"),
                      space='RGB', transparency=0)
  colormap<-data.frame(color_val=as.character(seq(-600, 600, 1)), color.name=as.character(col_fun(seq(-600, 600, 1))))
  
  dend <- dendrapply(dend, colorDendro, colormap, coloring_cond=coloring_cond, df_s=df_s)
  
  col <- c()
  for (item in res_cut){
    if (item %in% color2_clusters){
      color <- color2
    } else if (item %in% color1_clusters){
      color <- color1 
    } else if (item %in% color3_clusters){
      color <- color3
    } else if (item == 0){
      color <- '#FFFFFF'
    } else if (item%%2 == 1){
      color <- '#7F7F7F'
    } else if (item%%2 == 0){
      color <- '#CDCDCD'
    }
    col <- c(col, color)
  }
  
  leaf_labels <- label_clusters_dend(clusters_to_label=clusters_to_label, dend=dend, res_cut=res_cut)
  
  pdf(file=filename, height=4, width=2)
  dend %>%
    #rank_branches(dend) %>% 
    set(what=c('branches_lwd'), 0.05) %>%
    set_labels(leaf_labels) %>%
    set("labels_cex", 0.1) %>%
    plot(type='rectangle', horiz=TRUE, axes=FALSE)
  colored_bars(colors=col, dend=dend, sort_by_labels_order=FALSE, horiz=TRUE, y_shift=0.1, y_scale=0.07, rowLabels='')
  dev.off()
}


## 03.Plotting

df_s <- read.csv('rDEI.csv', row.names=1)

plot_dendrogram(dend=dend,
                df_s=df_s,
                coloring_cond='shortday.longday',
                color1_clusters=c(26),
                color2_clusters=c(25),
                color3_clusters=c(21),
                clusters_to_label=c(),
                filename=paste(c(pipe_out, '/fig_dend_', cond, '_col-sdld.pdf'), collapse=''))
```
