## Supplementary Codes for "A metabolic coincidence mechanism controls winter photoperiodism in plants": func_ClusterProfiler.R

```
library(enrichplot)
library(clusterProfiler)
library(org.At.tair.db)
library(biomaRt)


ensembl.arath <- useMart(biomart="plants_mart", host="plants.ensembl.org", dataset="athaliana_eg_gene")


call_enrichGO <- function(clusters, df_s, ont, threshold, out_dir, sim_cutoff=0.5){
  
  datasets <- listDatasets(useMart(biomart="plants_mart", host="plants.ensembl.org"))
  ensembl.arath = useMart(biomart="plants_mart", host="plants.ensembl.org", dataset="athaliana_eg_gene")
  
  for (m in 1:length(clusters)){
    
    print(m)
    
    probe_ids = strsplit(clusters[m], '\t')[[1]]
    entrez_ids = subset(df_s, probe_id %in% probe_ids)$entrez_id
    
    genes.q <- as.character(entrez_ids[!is.na(entrez_ids)])
    
    if (length(genes.q) < threshold){next}
    
    genes.b <- unlist(strsplit(as.character(df_s$entrez_id), "\t"))
    genes.b <- as.character(genes.b[!is.na(genes.b)])
    
    ego <- enrichGO(gene          = genes.q,
                    universe      = genes.b,
                    OrgDb         = org.At.tair.db,
                    ont           = ont,
                    pAdjustMethod = "BH",
                    pvalueCutoff  = 0.05,
                    qvalueCutoff  = 0.05,
                    readable      = FALSE,
                    keyType       = "ENTREZID")
    
    if (is.null(ego)==TRUE){next}
    if (dim(ego)[1]<1){next}
    
    #gene.df <- bitr(genes,
    #                fromType   = "TAIR",
    #                toType     = c("ENTREZID"),
    #                OrgDb      = org.At.tair.db)
    #genes[!genes %in% gene.df[,1]]
    
    egos <- simplify(ego, cutoff = sim_cutoff, by = "p.adjust", select_fun = min, measure = "Wang", semData = NULL)
    
    if (nrow(egos@result)>0){
      
      tair_loci <- c()
      tair_symbols <- c()
      
      for (i in 1:nrow(egos@result)){
        values <- egos@result$geneID[i]
        values <- strsplit(values, '/')
        
        if (length(values)>0){
          res <- getBM(attributes = c('tair_locus',
                                      'tair_symbol'), 
                       filters = 'entrezgene_id', 
                       values = values,
                       mart = ensembl.arath)
          tair_loci <- c(tair_loci, paste(res$tair_locus, collapse='\t'))
          tair_symbols <- c(tair_symbols, paste(res$tair_symbol, collapse='\t'))
          
        } else {
          tair_loci <- c(tair_loci, 'NA')
          tair_symbols <- c(tair_symbols, 'NA')
        }
        
      }
      
      egos@result$tair_identifier <- tair_loci
      egos@result$gene_symbol <- tair_symbols
    }
    
    cluster_lab <- paste(c(rep('0', 3-nchar(as.character(m))), as.character(m)), collapse='')

    dir.create(paste(c(out_dir), collapse=''), showWarnings = FALSE)
    dir.create(paste(c(out_dir, 'full/'), collapse=''), showWarnings = FALSE)
    dir.create(paste(c(out_dir, 'simplified/'), collapse=''), showWarnings = FALSE)

    write.csv(ego, paste(c(out_dir, "full/enrichGO_", ont, "_cluster", cluster_lab, ".csv"), collapse=''), row.names = FALSE)
    write.csv(egos, paste(c(out_dir, "simplified/enrichGO_", ont, "_cluster", cluster_lab, ".csv"), collapse=''), row.names = FALSE)
  }
}


call_enrichKEGG <- function(clusters, threshold, out_dir, df_s){
  
  datasets <- listDatasets(useMart(biomart="plants_mart", host="plants.ensembl.org"))
  ensembl.arath = useMart(biomart="plants_mart", host="plants.ensembl.org", dataset="athaliana_eg_gene")
  
  for (m in 1:length(clusters)){
    
    print(m)
    
    probe_ids = strsplit(clusters[m], '\t')[[1]]
    entrez_ids = subset(df_s, probe_id %in% probe_ids)$entrez_id
    
    genes.q <- as.character(entrez_ids[!is.na(entrez_ids)])

    if (length(genes.q) < threshold){next}
    
    genes.b <- unlist(strsplit(as.character(df_s$entrez_id), "\t"))
    genes.b <- as.character(genes.b[!genes.b=='NA'])
    
    ego <- enrichKEGG(gene        = genes.q,
                      universe      = genes.b,
                      organism      = 'ath',
                      pvalueCutoff  = 0.05,
                      keyType       = "ncbi-geneid")
    
    if (is.null(ego)==TRUE){next}
    if (dim(ego)[1]<1){next}
    
    if (nrow(ego@result)>0){
      tair_loci <- c()
      tair_symbols <- c()
      
      for (i in 1:nrow(ego@result)){
        values <- ego@result$geneID[i]
        values <- strsplit(values, '/')
        
        if (length(values)>0){
          res <- getBM(attributes = c('tair_locus',
                                      'tair_symbol'), 
                       filters = 'entrezgene_id', 
                       values = values,
                       mart = ensembl.arath)
          tair_loci <- c(tair_loci, paste(res$tair_locus, collapse='\t'))
          tair_symbols <- c(tair_symbols, paste(res$tair_symbol, collapse='\t'))
        } else {
          tair_loci <- c(tair_loci, 'NA')
          tair_symbols <- c(tair_symbols, 'NA')
        }
        
      }
      
      ego@result$tair_identifier <- tair_loci
      ego@result$gene_symbol <- tair_symbols
    }
    
    cluster_lab <- paste(c(rep('0', 3-nchar(as.character(m))), as.character(m)), collapse='')
    
    dir.create(out_dir, showWarnings = FALSE)
    write.csv(ego, paste(c(out_dir, "KEGG_cluster", cluster_lab, ".csv"), collapse=''), row.names = FALSE)
  }
}

## Calling the functions to perform enrichment of BP GO terms
mat_s <- read.csv("rDEI_ID.csv", stringsAsFactors=FALSE, dec='.')
call_enrichGO(df_s=mat_s, clusters=wg_clusters, ont='BP', out_dir=paste(c('BP1/'), collapse=''), threshold = 0, sim_cutoff = 1.0)
```
