## Supplementary Codes for "A metabolic coincidence mechanism controls winter photoperiodism in plants": func_dendextend.R

```
library(cluster)
library(dendextend)
library(scales)

order_clusters_by_median <- function(dend, res_DTC, cond){ 
  
  original_sequence <- res_DTC[order.dendrogram(dend)]
  probe_sequence <- original_sequence[!original_sequence==0]
  
  middle_probes <- c()
  for (n in unique(probe_sequence)){
    probes <- names(probe_sequence)[which(probe_sequence %in% n)]
    m_probe <- probes[length(probes)%/%2]
    middle_probes <- c(middle_probes, m_probe)
  }
  
  middle_probes_in_order <- c()
  for (probe in names(probe_sequence)){
    if (probe %in% middle_probes){
      middle_probes_in_order <- c(middle_probes_in_order, probe)
    }
  }
  
  clusters_in_order_new <- c()
  for (probe in middle_probes_in_order){
    old_cluster <- probe_sequence[[probe]]
    clusters_in_order_new <- c(clusters_in_order_new, old_cluster)
  }
  
  res_cut <- c()
  for (probe in original_sequence){
    if (probe==0){
      cluster_order <- 0
    } else{
      cluster_order <- match(probe, clusters_in_order_new)
    }
    res_cut <- c(res_cut, cluster_order)
  }
  names(res_cut) <- names(original_sequence)
  return(res_cut) 
}

make_cluster_descriptions <- function(clusters, df_s){

  ## Combine the information into a dataframe
  merged.cluster.list <- list()
  for (n in 1:length(clusters)){
    probe.list <- strsplit(clusters[n], '\t')[[1]]
    merged.probes <- clusters[n]
    merged.size <- length(probe.list)
    
    merged.FC.list <- list()
    merged.pVal.list <- list()
        
    conditions <- c('shortday.longday', 'shortday.LER_SD', 'LER_SD.shortday', 'lhyox_SD.LER_SD', 'phyB9_SD.COL_SD', 'lux.2_LDHH.COL_LDHH')

    for (p in 1:length(conditions)){
      FC.values <- df_s[rownames(df_s) %in% probe.list, conditions[p]]
      merged.FC <- 2**mean(log(FC.values, 2))
      merged.pVal <- wilcox.test(log(FC.values, 2), mu=0, alternative='two.sided', exact=NULL, correct=TRUE)$p.value
      merged.pVal <- merged.pVal*length(clusters)
      merged.FC.list[[p]] <- merged.FC
      merged.pVal.list[[p]] <- merged.pVal
    }
    
    merged.atID <- paste(df_s[rownames(df_s) %in% probe.list, 'locus_id'], collapse='\t')
    merged.eID <- paste(df_s[rownames(df_s) %in% probe.list, 'entrez_id'], collapse='\t')
    merged.name <- paste("cluster",as.character(n-1), sep='')
    #merge the probe list
    merged.cluster.list[[n]] <- c(merged.name, merged.size,
                                  merged.FC.list[[1]], merged.pVal.list[[1]],
                                  merged.FC.list[[2]], merged.pVal.list[[2]], 
                                  merged.FC.list[[3]], merged.pVal.list[[3]], 
                                  merged.FC.list[[4]], merged.pVal.list[[4]], 
                                  merged.FC.list[[5]], merged.pVal.list[[5]],
                                  merged.FC.list[[6]], merged.pVal.list[[6]], 
                                  merged.probes, merged.eID, merged.atID)
  }
  df_mi <- do.call("rbind", merged.cluster.list)
  colnames(df_mi) <- c('name', 'size',
                       'mean_rDEI_(shortday/longday)', 'p.adjust_(shortday/longday)',
                       'mean_rDEI_(shortday/LER_SD)', 'p.adjust_(shortday/LER_SD)', 
                       'mean_rDEI_(LER_SD/shortday)', 'p.adjust_(LER_SD/shortday)', 
                       'mean_rDEI_(lhyox_SD/LER_SD)', 'p.adjust_(lhyox_SD/LER_SD)', 
                       'mean_rDEI_(phyB9_SD/COL_SD)', 'p.adjust_(phyB9_SD/COL_SD)', 
                       'mean_rDEI_(lux-2_LDHH/COL_LDHH)', 'p.adjust_(lux-2_LDHH/COL_LDHH)', 
                       'probes', 'entrez ID', 'locus_id')
  
  return(df_mi)
}


colorDendro <- function(node, colormap, coloring_cond, df_s){
  probe <- get_leaves_attr(node, 'label')
  FC_sum <- mean(log(df_s[rownames(df_s) %in% probe, coloring_cond], 2))
  color_val <- as.character(round(FC_sum,2)*100)
  edgecol <- colormap[match(color_val, colormap$color_val),2]
  attr(node, "edgePar") <- list(col = as.character(edgecol))
  return(node)
}

convert_to_cluster_vector <- function(res){
  clusters <- c()
  for (n in min(res):max(res)){
    probes <- names(res[res==n])
    clusters <- c(clusters, paste(probes, collapse='\t'))
  }
  return(clusters)
}

label_clusters_dend <- function(clusters_to_label, dend, res_cut){
  leaf_labels <- rep(c(''), length(labels(dend)))
  for (n in clusters_to_label){
    probes <- names(res_cut)[which(res_cut %in% n)]
    m_probe <- probes[length(probes)%/%2]
    m_probe_ind <- match(m_probe, names(res_cut))
    leaf_labels[m_probe_ind] <- n
  }
  return(leaf_labels)
}
```
