## Supplementary Codes for "A metabolic coincidence mechanism controls winter photoperiodism in plants": README.txt

The 3 csv files: rDEI.csv, rDEI_ID.csv and diurnalValues.csv were used as input for the code.
diurnalValues.csv is a re-organised format of the data downloaded from the DIURNAL database website.
rDEI.csv and rDEI_ID.csv contains rDEI of genes calculated based on diurnalValues.csv.
