## Supplementary Codes for "A metabolic coincidence mechanism controls winter photoperiodism in plants": 1_checkClusterNumber.py

```
import pandas as pd
import numpy as np
import math
import matplotlib.pyplot as plt
from sklearn.cluster import KMeans
from sklearn.metrics import silhouette_samples, silhouette_score, pairwise_distances
import random
from scipy import stats


## Reading in tables
df_ts = pd.read_csv('diurnalValues.csv', index_col=0, header=0).sort_index()
df_s = pd.read_csv('rDEI.csv', index_col=0, header=0).sort_index()
df_lsdr = np.log2(df_ts.loc[:, 'longday_0':'shortday_44'])

## Z-score
df_lsdz = pd.DataFrame(stats.zscore(df_lsdr, axis=1, ddof=1), columns=df_lsdr.columns, index=df_lsdr.index)

## Select by rDEI
genes = df_s.loc[df_s['sd_ld']>2.0, :].index
X = df_lsdz.loc[genes, :]
range_n_clusters = [i for i in range(2, 10)]

## Choose optimal cluster number
inertia = []
silhouette_scores = []
for n_clusters in range_n_clusters:
    clusterer = KMeans(n_clusters=n_clusters, random_state=0).fit(X)
    inertia.append(clusterer.inertia_)
    silhouette_scores.append(silhouette_score(X, clusterer.fit_predict(X)))
    
fig, ax = plt.subplots(1, 2, figsize=(16, 4)) 
ax[0].plot(range_n_clusters, inertia)
ax[1].plot(range_n_clusters, silhouette_scores)
```
