## Supplementary Codes for "A metabolic coincidence mechanism controls winter photoperiodism in plants": 2_clustering.py

```
import pandas as pd
import numpy as np
import math
import matplotlib.pyplot as plt
from sklearn.cluster import KMeans
from sklearn.metrics import silhouette_samples, silhouette_score, pairwise_distances
import random
from scipy import stats

## Reading in tables
df_ts = pd.read_csv('diurnalValues.csv', index_col=0, header=0).sort_index()
df_s = pd.read_csv('rDEI.csv', index_col=0, header=0).sort_index()
df_lsdr = np.log2(df_ts.loc[:, 'longday_0':'shortday_44'])
df_lsdz = pd.DataFrame(stats.zscore(df_lsdr, axis=1, ddof=1), columns=df_lsdr.columns, index=df_lsdr.index)

## Selecting only genes with rDEI over 2.0
genes = df_s.loc[df_s['shortday/longday']>2.00, :].index
X = df_lsdz.loc[genes, :]

## Kmeans clustering
cluster_labels = KMeans(n_clusters=4, random_state=0).fit(X).fit_predict(X)
clusters = []

## Organising clustering results
for lab in np.unique(cluster_labels):
    cluster = list(np.compress(list(cluster_labels==lab), X.index))
    clusters.append(cluster)   
clusters.sort(key=len, reverse=True)

## Writing results to a text file
path = './'
with open(path+'clusters_kmeans_4_wg.txt', 'w') as file:
    for n in clusters:
        file.write('\t'.join(n)+'\n')
```
