## Supplementary Codes for "A metabolic coincidence mechanism controls winter photoperiodism in plants": 3_annotation.R

```
## GO term analysis
source('func_ClusterProfiler.R')

wg_clusters = readLines('clusters_kmeans_4_wg.txt')

mat_s <- read.csv("rDEI_ID.csv", stringsAsFactors=FALSE, dec='.')
path <- './'
call_enrichGO(df_s=mat_s, clusters=wg_clusters, ont='BP', out_dir=paste(c(path, 'BP/'), collapse=''), threshold = 0)
call_enrichGO(df_s=mat_s, clusters=wg_clusters, ont='CC', out_dir=paste(c(path, 'CC/'), collapse=''), threshold = 0)
call_enrichGO(df_s=mat_s, clusters=wg_clusters, ont='MF', out_dir=paste(c(path, 'MF/'), collapse=''), threshold = 0)
call_enrichKEGG(df_s=mat_s, clusters=wg_clusters, out_dir=paste(c(path, 'KEGG/'), collapse=''), threshold = 0)
```
